# A novel candidate neuromarker of central motor dysfunction in childhood apraxia of speech

**DOI:** 10.1101/2024.08.11.607491

**Authors:** Ioanna Anastasopoulou, Douglas O. Cheyne, Pascal van Lieshout, Peter H. Wilson, Kirrie J. Ballard, Blake W. Johnson

## Abstract

Childhood apraxia of speech (CAS) is conceived as an impairment of the central motor system’s ability to program multiple speech movements, resulting in inaccurate transitions between and relative timing across speech sounds. However, the extant neuroimaging evidence base is scant and inconclusive and the neurophysiological origins of these motor planning problems remain highly underspecified. In the first magnetoencephalography study of this disorder, we measured brain activity from typically developing children (N = 19) and children with CAS (N=7) during performance of a speech task designed to interrogate function of the speech areas of primary sensorimotor cortex. Relative to their typically developing peers, our sample of CAS children showed abnormal speech-related responses within the mu-band motor rhythm, and beamformer source reconstruction analyses specify a brain origin of this speech rhythm in the left cerebral hemisphere, within or near pre-Rolandic motor areas crucial for the planning and control of speech and oromotor movements. These results provide a new and specific candidate mechanism for the core praxic features of CAS; point to a novel and robust neurophysiological marker of typical and atypical expressive speech development; and support an emerging neuroscientific consensus which assigns a central role for programming and coordination of speech movements to the motor cortices of the precentral gyrus.

## Introduction

Childhood apraxia of speech (CAS) is a rare developmental motor speech disorder estimated to affect 0.1% - 0.2% of pre-schoolers (Shriberg et al., 1997). Individuals with CAS present with severe and highly persistent speech problems similar to those of adults with acquired apraxia of speech (AOS), characterized by inconsistent speech errors, lengthened and impaired co-articulatory transitions, and inappropriate prosody (ASHA, 2007). A key clinical feature of CAS is an absence of neuromuscular problems or weakness in the articulatory periphery (Ziegler & Ackermann, 2017). Accordingly, the core deficits of this disorder have been assigned an origin in the central nervous system, ostensibly within brain mechanisms associated with the ability to program and plan speech movements (ASHA, 2007; Morgan & Webster, 2018). Such processes would presumably be fundamental to the acquisition of skilled speech production in the typically developing human brain, developmental processes that remain largely and fundamentally unknown. Thus, insights into the neurophysiological and neuroanatomical origins of CAS have the potential to shed considerable light on the mechanisms that support and permit development of this crucial and uniquely human capacity.

Our current understanding of the brain origins of CAS remains limited by a small body of empirical neuroimaging and electrophysiological studies. To our knowledge the existing literature consists of 15 studies published over a period of about the last 25 years (Table 1). Their salient findings can be summarised as follows:

1. In contrast to most cases of acquired AOS, standard clinical MRI scans of individuals with CAS show no evidence of clinically significant lesions (Chilosi et al., 2015); or in a small proportion of cases, minor abnormalities with no clear relationship to CAS features (Chilosi et al., 2022);
2. Brain structure and function of genotyped members of the KE family with *FOXP2*-related CAS phenotype (Lai et al., 2001) have been characterised in a series of studies (Belton et al., 2003; Liégeois et al., 2003, 2011, 2016; Vargha-Khadem et al., 1998; Watkins et al., 2002). Relative to unaffected family members and unrelated controls, affected KE family members exhibit abnormal volumes in distributed cortical and subcortical structures. Functional neuroimaging has similarly shown abnormalities in distributed regions of premotor, supplementary, and primary motor cortices, cerebellum and basal ganglia.
3. It is unclear to what extent neuroimaging results from samples with FOXP2 mutations can be extrapolated to other neurogenetic CAS phenotypes or the CAS population. For example, while bilateral abnormalities of the basal ganglia are a consistent feature of the KE family, participants from a separate multigenerational CAS family did not show any significant basal ganglia abnormalities (Liégeois t al., 2019). Complicating this picture further is the fact that only a small proportion of CAS cases present with a clear neurogenetic basis (Morgan & Webster, 2018).
4. EEG studies have reported abnormal auditory mismatch responses to phonemic contrasts in speech stimuli (Froud & Khamis-Dakwar, 2012); and abnormal speech movement-related brain potentials (Preston et al., 2014).
5. Therapeutic interventions have been reported to result in significant volume changes in both cortical gray matter (Kadis et al., 2014) and white matter fibre tracts (Fiori et al., 2021).

**Table 1.**
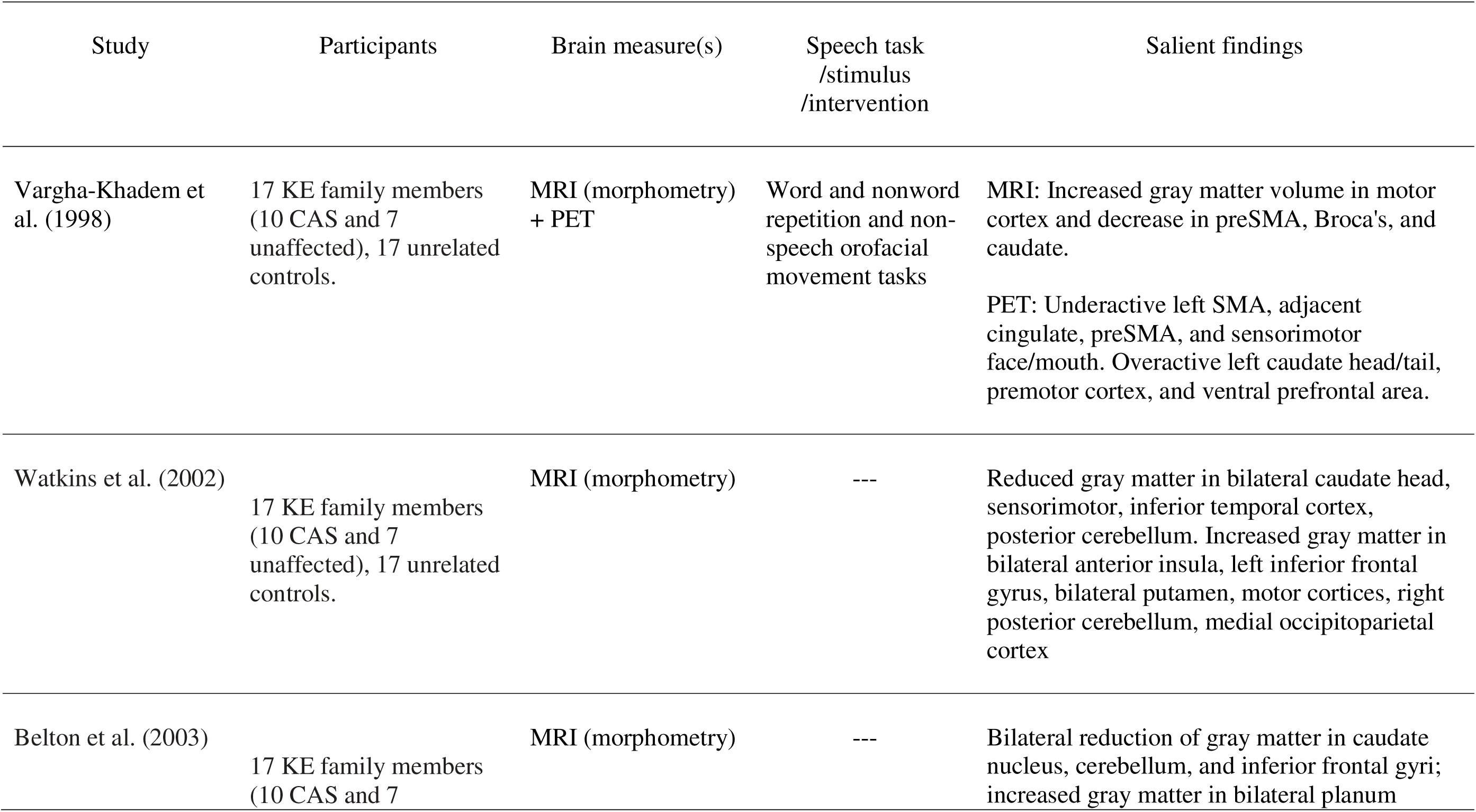
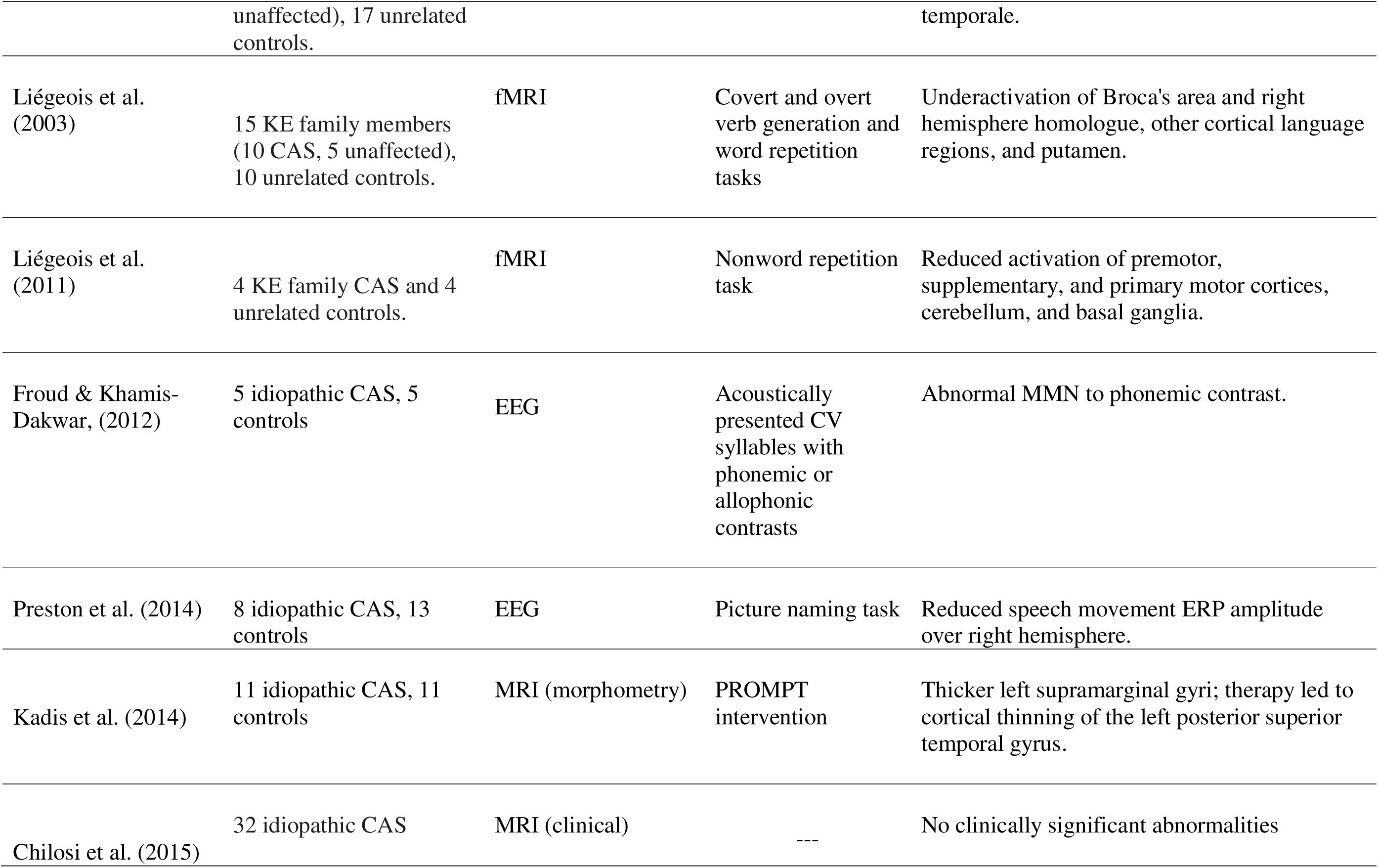
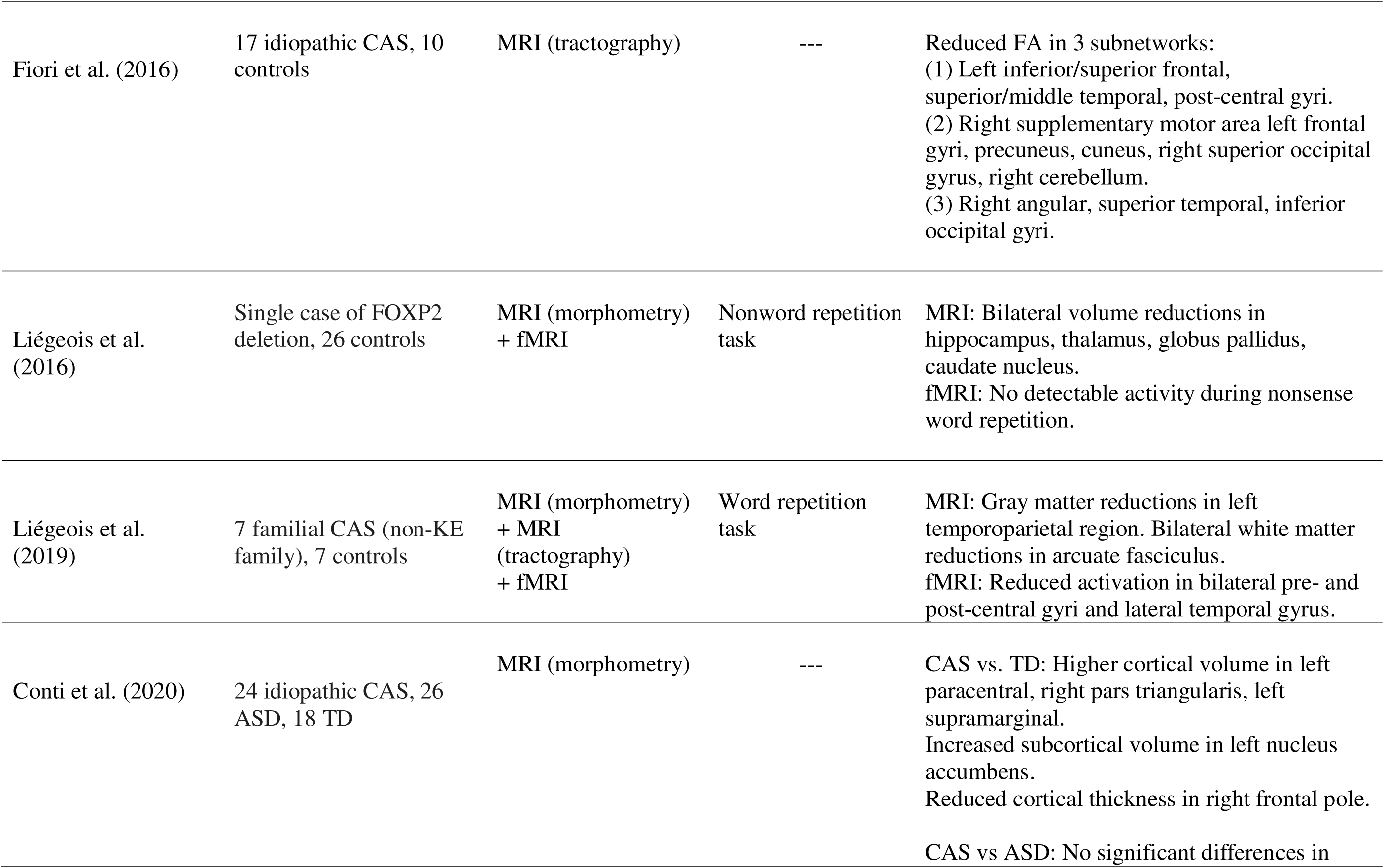
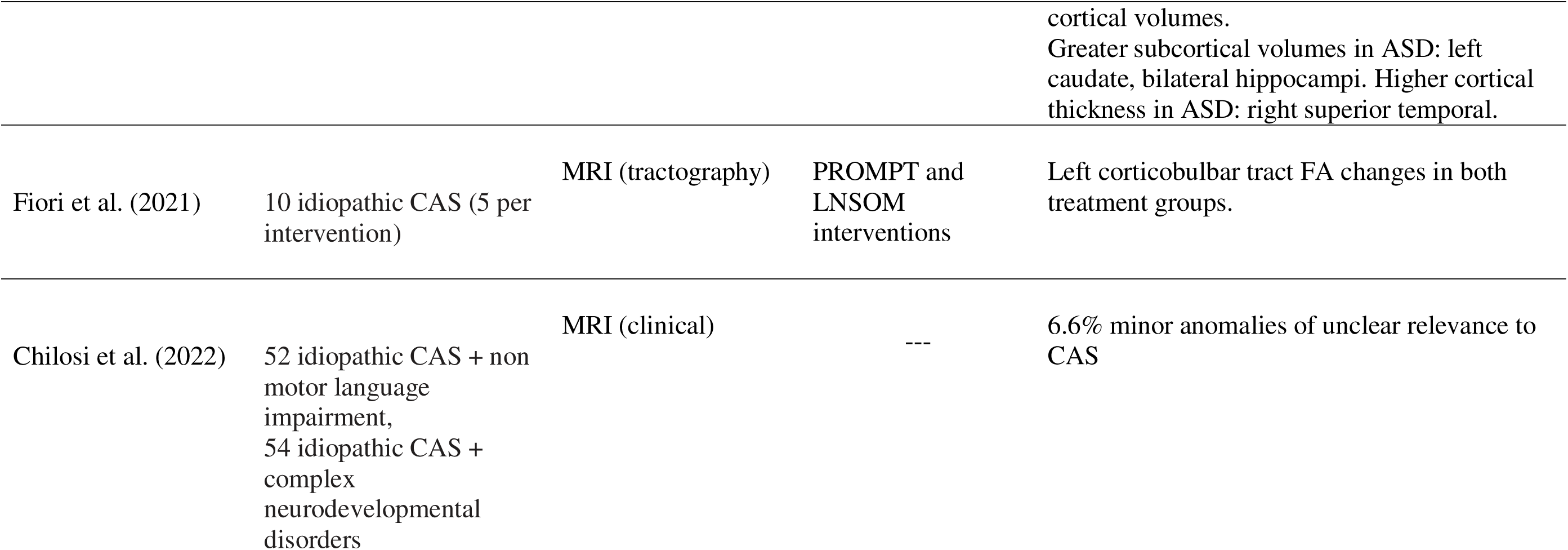
Neuroimaging and neurophysiological studies of childhood apraxia of speech.

| Study | Participants | Brain measure(s) | Speech task<br>/stimulus<br>/intervention | Salient findings |
| --- | --- | --- | --- | --- |
| Vargha-Khadem et al. (1998) | 17 KE family members (10 CAS and 7 unaffected), 17 unrelated controls. | MRI (morphometry) + PET | Word and nonword repetition and non-speech orofacial movement tasks | MRI: Increased gray matter volume in motor cortex and decrease in preSMA, Broca's, and caudate.<br>PET: Underactive left SMA, adjacent cingulate, preSMA, and sensorimotor face/mouth. Overactive left caudate head/tail, premotor cortex, and ventral prefrontal area. |
| Watkins et al. (2002) | 17 KE family members (10 CAS and 7 unaffected), 17 unrelated controls. | MRI (morphometry) | --- | Reduced gray matter in bilateral caudate head, sensorimotor, inferior temporal cortex, posterior cerebellum. Increased gray matter in bilateral anterior insula, left inferior frontal gyrus, bilateral putamen, motor cortices, right posterior cerebellum, medial occipitoparietal cortex |
| Belton et al. (2003) | 17 KE family members (10 CAS and 7 | MRI (morphometry) | --- | Bilateral reduction of gray matter in caudate nucleus, cerebellum, and inferior frontal gyri; increased gray matter in bilateral planum |
| Speech motor cortex function in CAS |  |  |  |  |
|  | unaffected), 17 unrelated controls. |  |  | temporale. |
| Liégeois et al. (2003) | 15 KE family members (10 CAS, 5 unaffected), 10 unrelated controls. | fMRI | Covert and overt verb generation and word repetition tasks | Underactivation of Broca's area and right hemisphere homologue, other cortical language regions, and putamen. |
| Liégeois et al. (2011) | 4 KE family CAS and 4 unrelated controls. | fMRI | Nonword repetition task | Reduced activation of premotor, supplementary, and primary motor cortices, cerebellum, and basal ganglia. |
| Froud & Khamis-Dakwar, (2012) | 5 idiopathic CAS, 5 controls | EEG | Acoustically presented CV syllables with phonemic or allophonic contrasts | Abnormal MMN to phonemic contrast. |
| Preston et al. (2014) | 8 idiopathic CAS, 13 controls | EEG | Picture naming task | Reduced speech movement ERP amplitude over right hemisphere. |
| Kadis et al. (2014) | 11 idiopathic CAS, 11 controls | MRI (morphometry) | PROMPT intervention | Thicker left supramarginal gyri; therapy led to cortical thinning of the left posterior superior temporal gyrus. |
| Chilosi et al. (2015) | 32 idiopathic CAS | MRI (clinical) | --- | No clinically significant abnormalities |
| Fiori et al. (2016) | 17 idiopathic CAS, 10 controls | MRI (tractography) | --- | Reduced FA in 3 subnetworks:<br>(1) Left inferior/superior frontal, superior/middle temporal, post-central gyri.<br>(2) Right supplementary motor area left frontal gyri, precuneus, cuneus, right superior occipital gyrus, right cerebellum.<br>(3) Right angular, superior temporal, inferior occipital gyri. |
| Liégeois et al. (2016) | Single case of FOXP2 deletion, 26 controls | MRI (morphometry) + fMRI | Nonword repetition task | MRI: Bilateral volume reductions in hippocampus, thalamus, globus pallidus, caudate nucleus.<br>fMRI: No detectable activity during nonsense word repetition. |
| Liégeois et al. (2019) | 7 familial CAS (non-KE family), 7 controls | MRI (morphometry) + MRI (tractography) + fMRI | Word repetition task | MRI: Gray matter reductions in left temporoparietal region. Bilateral white matter reductions in arcuate fasciculus.<br>fMRI: Reduced activation in bilateral pre- and post-central gyri and lateral temporal gyrus. |
| Conti et al. (2020) | 24 idiopathic CAS, 26 ASD, 18 TD | MRI (morphometry) | --- | CAS vs. TD: Higher cortical volume in left paracentral, right pars triangularis, left supramarginal.<br>Increased subcortical volume in left nucleus accumbens.<br>Reduced cortical thickness in right frontal pole.<br>CAS vs ASD: No significant differences in cortical volumes.<br>Greater subcortical volumes in ASD: left caudate, bilateral hippocampi. Higher cortical thickness in ASD: right superior temporal. |
| Fiori et al. (2021) | 10 idiopathic CAS (5 per intervention) | MRI (tractography) | PROMPT and LNSOM interventions | Left corticobulbar tract FA changes in both treatment groups. |
| Chilosi et al. (2022) | 52 idiopathic CAS + non motor language impairment,<br>54 idiopathic CAS + complex neurodevelopmental disorders | MRI (clinical) | --- | 6.6% minor anomalies of unclear relevance to CAS |

As it currently stands, this body of research constitutes solid support for both structural and functional anomalies within broadly defined speech and language brain networks, but provides little further specificity for mechanisms that may directly contribute to the core praxic features of CAS. The present study contributes to this literature with the first magnetoencephalographic (MEG) neuroimaging data from a clinically well-defined sample of children with idiopathic CAS, and typically developing controls. We aimed to constrain the current highly underspecified picture of CAS mechanisms by interrogating function within speech centres in the proximity of primary speech motor cortex (Anastasopoulou et al., 2024; Riecker et al., 2000), using a speech task with minimal requirements for linguistic and attentional/memorial processing (Frankford et al., 2021) and an emphasis on sensorimotor phonological and phonetic sequencing operations (Riecker et al., 2000).

## Materials and Methods

### Participants and group assignment

Nineteen typically developing (TD) children (8 females, mean age =11.0 years; SD = 2.5, range 7.5 – 16.7) and seven children with CAS (7 males, mean age = 8.9 years, SD = 2.2, range = 6.8 - 12.8) participated in the study. For statistical analyses of demographic and clinical data, seven individuals of the TD group (AM group) were age-matched to the individuals of the CAS group. All children were right-handed as assessed by a short version of the Edinburgh handedness questionnaire (Veale, 2014). All procedures were approved by the Macquarie University Human Subjects Research Ethics Committee.

To provide a further developmental contrast for the TD children, spectro-temporal analyses were carried out in a reanalysis of data acquired in a separate experiment (Anastasopoulou et al., 2024) from a group of 10 adult participants (4F; mean age 32.5, range 19.7–61.8; all right-handed) and one additional adult participant (M, 63.2 years) not included in the dataset described in (Anastasopoulou et al., 2024). All experimental tasks and procedures for the adult (AD) group were identical to those described below for the children, with the exceptions that (1) structural brain scans were obtained (and co-registered with MEG for source reconstruction) for adults but not child participants; and (2) only the child groups participated in speech and motor assessments described in the following.

### CAS diagnosis

A CAS diagnosis required (a) the three features established by consensus in the ASHA Technical Report (2007) and (b) a minimum of four out of the 10 features outlined in Strand’s 10-point checklist (Murray et al., 2015; Shriberg et al., 2009) across at least three assessment tasks. Two certified speech-language pathologists (authors I.A. and K.B.) independently reviewed and rated audio-video recordings of speech productions obtained during administration of the Single-Word Test of Polysyllables (Gozzard et al., 2008). Of ten children with suspected CAS initially enrolled in the study, two children who did not meet the diagnostic criteria for CAS, and one left-handed child were excluded from further analysis. Interrater reliability for CAS diagnosis was 87.5%.,

### Speech and motor assessments

All children were screened for pure-tone hearing thresholds and completed a battery of speech, expressive and receptive language, and gross and fine motor skill assessments: The Sounds-in-Words subtest of the Goldman-Fristoe Test of Articulation–Third Edition (GFTA-3; (Goldman & Fristoe, 2015); receptive and expressive language components of the Clinical Evaluation of Language Fundamentals test (CELF-5; (Wiig et al., 2013); Verbal Motor Production Assessment for Children (VMPAC-R, (Hayden & Namasivayam, 2021); and the Movement Assessment Battery for Children (ABC-2, (Henderson et al., 2007). Caregivers completed developmental coordination (DCDQ, (Wilson et al., 2009)) and handedness questionnaires (Veale, 2014).

### MEG recordings

Neuromagnetic brain activity was recorded with a KIT-Macquarie MEG160 (Model PQ1160R-N2, KIT, Kanazawa, Japan) whole-head MEG system consisting of 160 first-order axial gradiometers with a 50-mm baseline (Kado et al., 1999; Uehara et al., 2003). MEG data were acquired with analogue filter settings of 0.3 Hz high-pass, 200 Hz low-pass, 1000 Hz sampling rate and 16-bit quantization. Measurements were carried out with participants in supine position in a magnetically shielded room (Fujihara Co. Ltd., Tokyo, Japan). Five head position indicator coils (HPI) were attached to an elastic cap on the head. HPI positions were measured at the beginning and at the end of the experiment, with a maximum displacement criterion of < 5 mm in any direction.

### Head shape templates

Individual structural MRI scans were not available for the child participants in this study so a surrogate MRI approach was used which warps a template brain to each subject’s digitized head shape using the iterative closest point algorithm implemented in SPM8 (Litvak et al., 2011) and the template scalp surface extracted with the FSL toolbox (Jenkinson et al., 2012); see also (Cheyne et al., 2014; Johnson et al., 2020) for application with child participants. Participant’s head shapes and fiducial locations were digitized prior to MEG recordings with a stylus digitiser (Polhemus FastTrack, Colchester, VT). For adult participants, MEG data were co-registered with individual T1-weighted anatomical magnetic resonance images (MRIs) acquired in a separate scanning session using a 3T Siemens Magnetom Verio scanner with a 12-channel head coil.

### Audio speech recordings

Time-aligned audio speech recordings were recorded in an auxiliary channel of the MEG setup with the same sample rate (1000 Hz) as the MEG recordings and were used to identify speech onset/offset events for use in MEG source reconstruction analyses.

An additional high fidelity speech recording was simultaneously recorded with an optical microphone (Optoacoustics, Or-Yehuda, Israel) fixed on the MEG dewar at a distance of 20 cm away from the mouth of the speaker; and digitised with a sound card at 48 kHz sample rate and 24-bit quantization precision. All participants were also fitted with four MEG-compatible speech movement tracking coils (Alves et al., 2016) placed on the upper and lower lips, tongue body, and jaw (Anastasopoulou et al., 2022, 2024). Analyses of the speech tracking and high-fidelity acoustic data will be presented in a separate report and are not further discussed here.

### Experimental design

The overall experiment consisted of a speech condition and a manual button press condition. For the speech condition, participants performed a *reiterated nonword speech task* which limits requirements for semantic, syntactic and attentional processing (Frankford et al., 2021) and emphasises motoric sequencing operations (Riecker et al., 2000). Previous fMRI (Riecker et al., 2000) and MEG (Anastasopoulou et al., 2024) studies of healthy adults have shown that the reiterated nonword task elicits spatially focal brain activations restricted to motor speech centres in or near the peri-Rolandic sensorimotor cortices. (In contrast, commonly used expressive speech mapping protocols such as sentence reading, semantic word judgements, picture naming, verb generation necessarily invoke memorial and cognitive/linguistic operations in addition to phonological, phonetic and sensorimotor operations, and these tasks typically elicit widely distributed activations in broad regions of prefrontal, temporal and parietal cortex; see (Agarwal et al., 2019; Munding et al., 2016) for reviews of fMRI and MEG expressive speech mapping studies.)

The speech task protocol is illustrated in Figure 1A. Following previously published protocols (Anastasopoulou et al., 2022, 2024; van Lieshout et al., 2007), participants were required to produce utterances of the disyllabic V1CV2 sequences /ipa/ and /api/ at a constant rate during the course of a single exhalation of a deep breath intake. Two speech rate conditions (normal and faster) were used for each production, for a total of four speech conditions: /ipa/ normal rate, /ipa/ faster rate, /api/ normal rate, and /api/ faster rate. Each participant generated ten “trial sets” (van Lieshout et al., 2007) for each speech condition, with each trial set lasting approximately 12 seconds. For each trial set, participants were instructed to take a deep breath and, for the normal rate production, to reiteratively utter the non-words in a comfortable, conversational rate during the course of a single breath exhalation. For the faster rate, they were instructed to produce the nonwords at a faster rate while maintaining accuracy (van Lieshout et al., 2002). Each 12 sec trial set resulted in about 10 separate nonword utterances at the normal rate and about 15 at the faster rate. An inter-trial set rest interval of 4 secs was terminated by instructions (1 sec) for fixation and breath intake for the next trial set. A short break was provided after performance of 10 trials sets of the same speech condition. Participants were instructed to minimize head movement and avoid blinking during speech trial sets.

**Figure 1.**
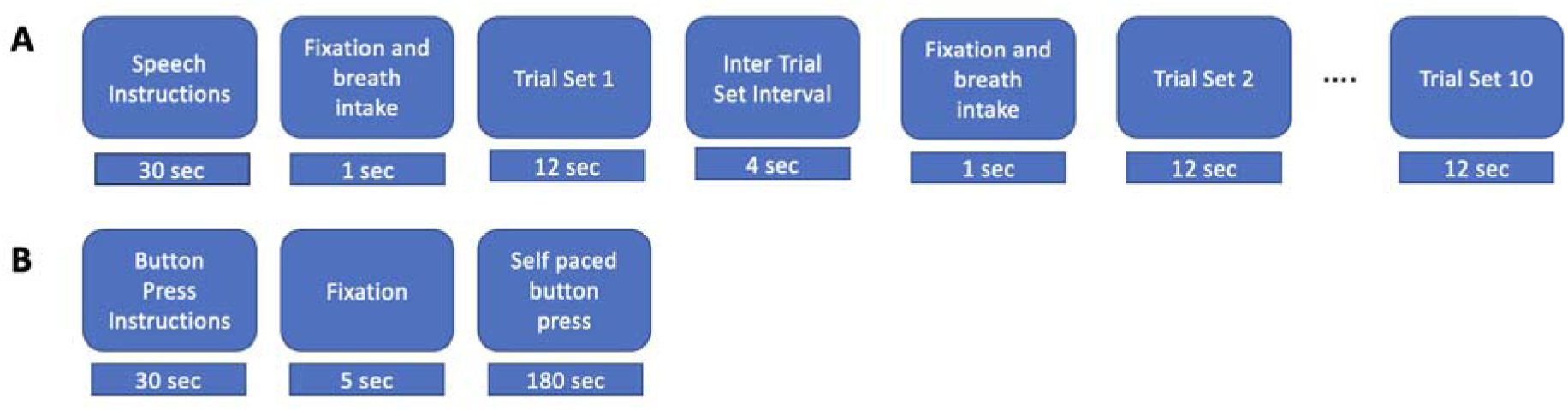
Experimental procedures. **A.** Speech task. Instructions were displayed for 30s, followed by an intertrial interval lasted for 15s and 5s fixation cross and breath intake in preparation for the speech production trial set. During a trial set participants produced the indicated nonword in a reiterated fashion for 12s. 10 consecutive trial sets were performed for each nonword stimulus. **B.** Button press task. Instructions were displayed for about 30s followed by a fixation cross, during which participants performed self-paced button pressed with the index finger of their dominant (right) hand at a rate of about 1 per 2 seconds for a total of about 90 trials.

For the manual task (Figure 1B), participants performed a button press task on a fibre optic response pad (Current Designs, Philadelphia) with the index finger of the dominant (right) hand at a self-paced rate of about 1 per 2 s for 180 s, resulting in a total of about 90 button press trials. In previous studies of manual motor control this task has been shown to elicit robust neuromagnetic responses that can localised in a straightforward manner to the hand region of primary sensorimotor cortex (e.g. Cheyne et al., 2014; De Nil et al., 2021; Johnson et al., 2020). This well-established response provides a useful functional-anatomic reference landmark for the results of the MEG speech analyses (Anastasopoulou et al., 2024).

### MEG analyses

#### A. Source reconstruction

##### Speech task

Speech trial set onsets were identified and marked from the speech channel of the MEG recordings. MEG data were segmented with an epoch of −10 sec to + 5 sec from the onset of each trial set, selected to encompass the final 5 sec of the previous trial set, the 5 sec inter-trial set rest period, and the first five sec of the current trial set (speech – rest – speech). Preliminary source analyses showed no significant difference in source location for different productions (/ipa/ versus /api/) or speech rates (faster versus slower). Accordingly, MEG data from all four speech conditions were averaged to maximize the signal-to-noise ratio, for a total of 40 trial sets/averaged epoch (4 speech conditions * 10 trial sets). All epoched data were digitally filtered with a bandpass of 0.3-100 Hz and a 50 Hz notch filter.

Source reconstruction was performed using the scalar synthetic aperture magnetometry (SAM) beamformer implemented in the BrainWave MATLAB toolbox (Jobst et al., 2018). The SAM noise-normalized (pseudo-t) beamformer images were computed using a frequency range of 18-22 Hz (center of the beta frequency band), a sliding active window of 0 to 1.0 sec (first second of current speech trial set) and a baseline window of −5 to −3 sec (first two seconds of inter trial set rest period) over 10 steps with a step size of 0.2 sec and pseudo-z beamformer normalisation (3 ft/sqrt (Hz) RMS noise). While beamforming analyses typically employ equal baseline and active window durations, in the present analyses we wished to use a longer baseline epoch to reduce the chance of a biased estimate of baseline power, which we reasoned is more likely to vary over time than during the active speech period (since reiterated speech is akin to a steady state). While larger order mismatches (e.g., comparing 1.0 s to 0.1 s) are likely to be problematic, simulation studies (Brookes et al., 2008) have shown that covariance errors are minimized, and therefore beamformer weights are stable, with at least 5 s of total data (a requirement that is well exceeded in the present case) and our preliminary analyses confirmed that a 2 s baseline did not affect the beamforming results relative to a 1 s baseline. The combined active and control windows were used to compute the data covariance matrix for beamformer weight calculations, while the full 15 s time window was used to compute data covariance and beamformer weights to extract single-trial time courses of localized neural activity (“virtual sensors”) for each trial.

Statistical analysis of group beamformer images was performed with cluster-based permutation testing (alpha = 0.05, 512-2048 permutations, omnibus correction for multiple comparisons).

Source reconstruction results of adult group speech and button press responses have been previously described by Anastasopoulou et al. (2024) in a study using identical experimental methods and procedures to those described here for the child participants.

### Manual task

MEG data were segmented into 1.5 s epochs comprising −0.5 sec to +1.0 sec with respect to button-press onset, resulting in about 90 button press trials. Epoched data sets were digitally filtered from 0.3-100 Hz and a 50 Hz notch filter. SAM source images were computed using a frequency range of 15-25 Hz (beta band), a sliding active window of 0.6 to 0.8 sec and a baseline window of −0.5 to −0.3 sec over 10 steps with a step size of 0.01 sec and pseudo-z beamformer normalisation (3 ft/sqrt (Hz) RMS noise). These time windows were chosen to encompass the known periods of maximal event-related synchronisation (ERS) and event-related desynchronisation (ERD) respectively of MEG motor rhythms in a button press task (see for e.g., (Cheyne et al., 2014; Johnson et al., 2020). Statistical analysis of group beamformer images was performed with cluster-based permutation testing (2048 permutations, omnibus correction for multiple comparisons).

#### B. Spectro-temporal analyses

##### Speech task

Time frequency plots were computed from virtual sensor locations at the centre of the mean beamformer map clusters obtained as described above. For the speech task, the AD and CAS group showed single clusters in the left and right hemispheres respectively (Figure 2). In order to assess hemispheric lateralisation of brain responses, the opposite hemisphere coordinates were obtained by reversing the sign of the X coordinate of the obtained cluster.

**Figure 2.**
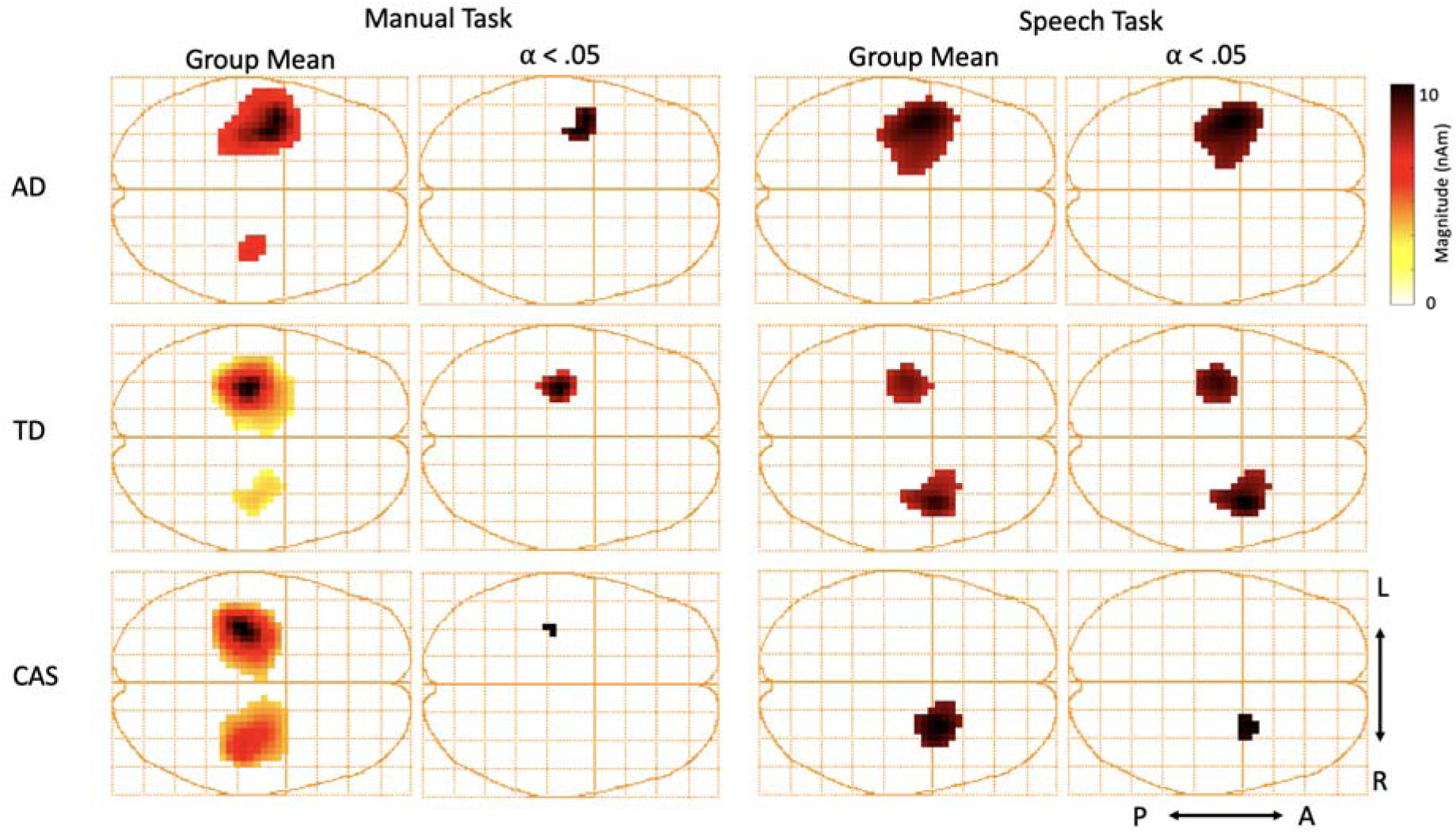
Group mean SAM beamformer maps (left panel) and group statistical maps (right panel) for manual and speech tasks. Statistical maps show cluster-based permutation analysis of SAM-beamformer source maps, cluster alpha = 0.05, 512-1024 permutations, omnibus correction for multiple comparisons. Axial view: L = left, R = right, A = anterior, P = posterior.

Time frequency plots were generated with a time range of −10 to +5 sec from the trial set onset and a frequency range of 1-100 Hz. Time-frequency data for each individual were subsequently partitioned and frequency averaged over three frequency bands: beta (15-25 Hz), mu (8-12 Hz) and theta (4-7 Hz). Based on the group-averaged time-frequency spectrograms, event-related synchronisation (ERS) and desynchronisation (ERD) time-epochs were chosen as −5 to −3 sec and −1 to 1 sec, respectively for beta and mu bands, and −1 to 1 sec and −5 to −3 sec respectively for theta band.

##### Manual task

Voxel locations at the centre of group mean clusters were used to generate virtual sensor time frequency plots with a time range of −0.5 to +1.0 sec from the button press onset and a frequency range of 1-100 Hz. Based on the group-averaged time-frequency spectrograms, event-related synchronisation (ERS) and desynchronisation (ERD) time-epochs were chosen as −0.2 to 0 sec, and 0.5 to 0.7 sec respectively, for beta and mu, and 0.5 to 0.7 sec and 0.2 to 0 sec respectively, for theta band.

##### Response Magnitude

For the purposes of the statistical analyses described below, a metric of “response magnitude” was defined as the average magnitude over the ERS epoch minus the average magnitude over the ERD epoch.

### Statistical analyses

1. <u>Between-group differences in response magnitude</u>. For each task (button press, speech), hemisphere (left and right) and frequency band (theta, mu, beta), Kruskal-Wallis non-parametric one-way ANOVA (MatLab *kruskalwallis* function) was computed to determine whether there was a significant group difference in median response magnitudes using a critical alpha = .05. Where a significant group difference was indicated by the results of the overall ANOVA, planned comparisons (CAS versus TD; TD versus AD) were performed using the nonparametric Wilcoxon rank sum test (MatLab *ranksum* function) for equal medians), using the Bonferroni correction for multiple comparisons. Where a significant CAS:TD group difference was obtained, a further Wilcoxon rank sum comparison between CAS and AM groups was performed was performed to assess whether the CAS:TD group difference still held when members of the TD group were age-matched to the CAS group.
2. <u>Within-group hemispheric differences in response magnitude</u>. For each group (CAS, TD, AD) and frequency band (theta, mu, beta), hemispheric differences were tested with nonparametric Wilcoxon signed rank test for zero median.
3. <u>Between-group differences in hemispheric lateralisation</u>. For each participant, a laterality index (LI) was computed from response magnitudes in left (LH) and right hemispheres (RH), according to the formula (LH-RH)/(LH+RH). LI values range from 1 to −1, where 1 indicates fully left-lateralised, −1 indicates fully right-lateralised, and 0 indicates fully bilateral. Statistical analyses were then performed as described for response magnitude above.
4. <u>Effects of age within the TD group</u>. For each task (button press, speech), hemisphere (left and right) and frequency band (theta, mu, beta), a linear regression model (MatLab *fitlm* function) was used to test for linear effects of age within the TD group.

## Results

### Speech and motor assessments

A summary of demographic and test results for TD, CAS, and AM groups are presented in Table 2. Test results for individual participants are provided in the supplementary materials.

**Table 2.**
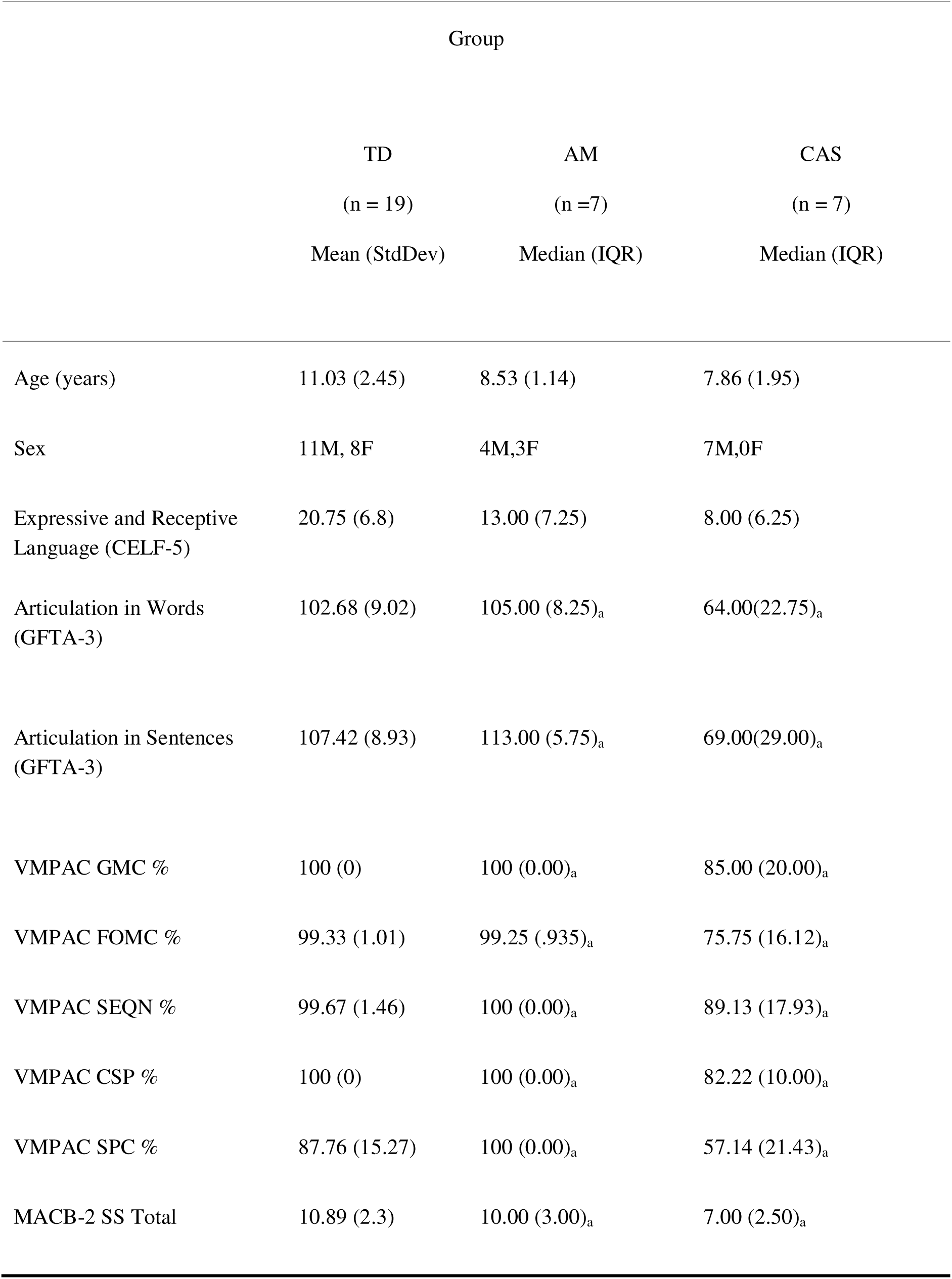
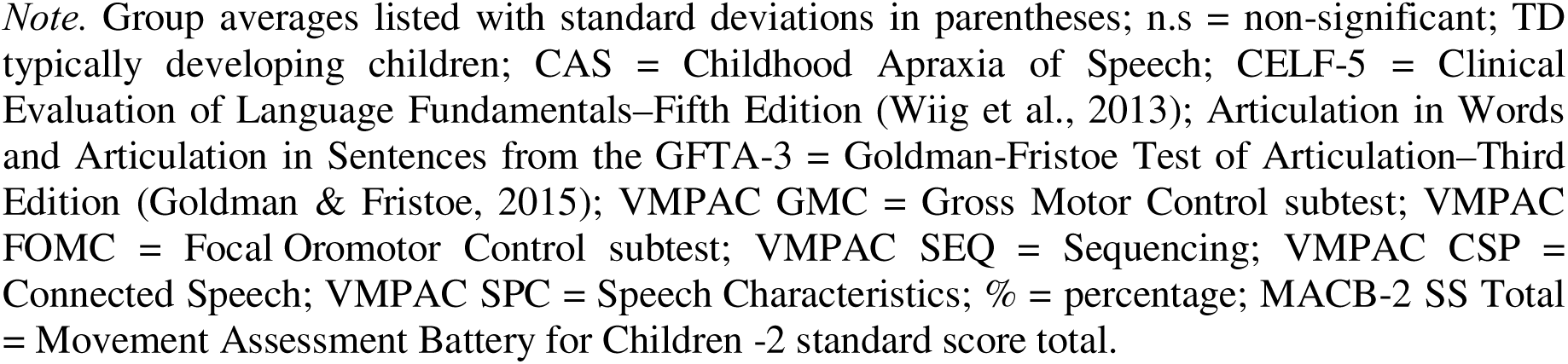
Demographic data and clinical test scores for child participants.

Statistical comparisons between the CAS group and AM group were carried out with nonparametric Wilcoxon signed rank tests (2-sided). Results showed no significant group difference in language abilities (CELF-5; *p* = .197), but CAS children performed significantly worse than age-matched TD on all measures of speech articulation (GFTA-3 Words, *p* = .001, GFTA Sentences, *p* = .001) and verbal motor production (VMPAC GMC%, *p* = .005; VMPAC FOMC%, *p* = .001; VMPAC SEQN%, *p* = .002; VMPAC CSP%, *p* = .001; VMPAC SPC%, *p* = .001). Further, CAS children also performed significantly worse on the measure of general gross and fine motor skills (MACB-2, *p* = .019).

In summary, the test results show that our sample of CAS children exhibited a typical CAS profile of specific and marked deficits in measures of articulation and verbal motor production within a normal and age-appropriate repertoire of non-speech language abilities. Further, our CAS sample also showed significant deficits in non-speech motor skills suggestive of probable developmental coordination disorder (DCD), in agreement with previous reports that have indicated a strong association between CAS and DCD (And & Dodd, 1996; Duchow et al., 2019; Iuzzini-Seigel, 2021; Iuzzini-Seigel et al., 2022; Knežević, 2019; McCabe et al., 1998).

### MEG data

#### a. Speech task

##### Source reconstruction

Group mean and corresponding statistical maps of SAM-beamformer images for the speech condition are shown in Figure 2 (right panels) and cluster coordinates are provided in Table 3. Source reconstruction indicates strikingly contrasting group results for the speech task. While source clusters for all three groups are aligned near the coronal midline (adjacent to the Rolandic fissure), adults show a single cluster in the left hemisphere, TD clusters are bilateral, while the CAS group shows a single significant cluster in the right hemisphere.

**Table 3.** Virtual sensor Talairach coordinates.

| **Manual task** |  |  |  |  |  |  |  |  |
| --- | --- | --- | --- | --- | --- | --- | --- | --- |
| **Left Hemisphere** |  |  |  |  | **Right Hemisphere** |  |  |  |
| **Group** | **X** | **Y** | **Z** | **Brain Region** | **X** | **Y** | **Z** | **Brain Region** |
| **AD** | -33 | -12 | 66 | Precentral gyrus BA4 | 38 | -16 | 65 | Precentral gyrus BA4 |
| **TD** | -30 | -17 | 43 | Precentral gyrus BA4 | 38 | -14 | 38 | Precentral gyrus BA4 |
| **CAS** | -30 | -21 | 51 | Precentral gyrus BA4 | 42 | -21 | 47 | Postcentral gyrus BA3 |

| **Speech task** |  |  |  |  |  |  |  |  |
| --- | --- | --- | --- | --- | --- | --- | --- | --- |
| **Left Hemisphere** |  |  |  |  | **Right Hemisphere** |  |  |  |
| **Group** | **X** | **Y** | **Z** | **Brain Region** | **X** | **Y** | **Z** | **Brain Region** |
| **AD** | -38 | -1 | 50 | Middle frontal gyrus BA6 | 38 | -1 | 50 | Middle frontal gyrus BA6 |
| **TD** | -30 | -10 | 24 | Insula BA 13 | 38 | 1 | 28 | Precentral gyrus BA6 |
| **CAS** | -22 | 20 | 19 | Subgyral | 22 | 20 | 19 | Subgyral |

*Note.* Coordinates correspond to the centre of group mean beamformer map clusters shown in Figure 2. For the unilateral clusters of AD and CAS groups for the speech task, the contralateral location was obtained by reversing the sign of the X coordinate of the obtained cluster.

##### Virtual sensors

Group mean virtual sensor time-frequency plots were generated from the speech task coordinates listed in Table 3 and are plotted in the left panels of Figure 3. The following features can be observed in these source-derived temporal spectrograms: (1) Substantial high-frequency broadband noise during the speech trial sets, attributable to myogenic activity that is an inevitable artifact of the overt speech task; (2) prominent mu/beta band (about 10-25 Hz) desynchronisation that is continuous during the speech trial sets, and also for about 2 seconds prior to the speech trial set onset, reflecting motor preparatory activity that is characteristic of the mu/beta bands (Cheyne, 2013). For adults, mean beta/mu band magnitude is notably lower in the right relative to the left hemisphere, is lower in magnitude for TD relative to adults, and is yet lower in magnitude for CAS relative to TD. (2) Theta band (circa 3-7 Hz) synchronisation which roughly tracks the time course of the mu/beta desynchronisation. (For aid in interpretation we note that the speech spectrograms are baselined to the first two seconds of the inter-trial set rest period, −5 to −3 sec. This baseline affects the visual appearance of the spectrogram during the immediately preceding time period, where the high-frequency noise and mu/beta-band desynchronisation appear to stop about 1 sec before the end of the speech trial set).

**Figure 3.**
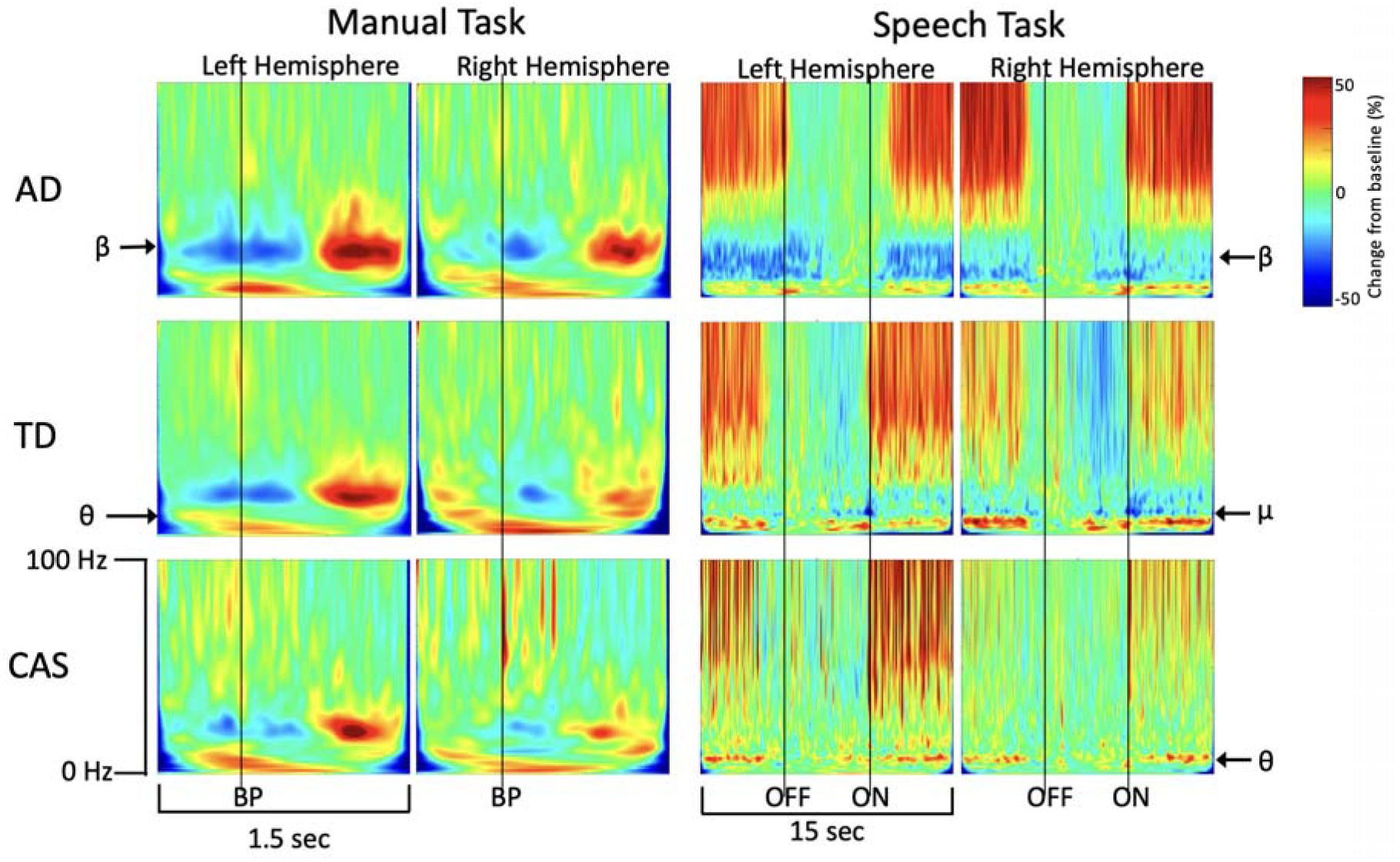
Group mean virtual sensor time-frequency spectrograms. All virtual sensor plots correspond to the Talairach coordinates listed in Table 3. Manual task data are baselined to the full epoch; speech task data are baselined −5 to −3 sec to emphasise beta band desynchronisation. BP = button press onset, OFF = speech trial set offset, ON = speech trial set onset.

Group mean comparisons of frequency-band averaged time series (Figure 4) further confirm that the three groups exhibit distinct patterns of brain activity during speech movements. For the beta and mu bands, adults show strikingly larger ERS magnitudes (during the non-speaking inter trials set rest period) than TD in both hemispheres, although TD do show speech-offset related ERS in the right hemisphere. No comparably distinct ERS or ERD epochs are apparent in the CAS average waveforms.

**Figure 4.**
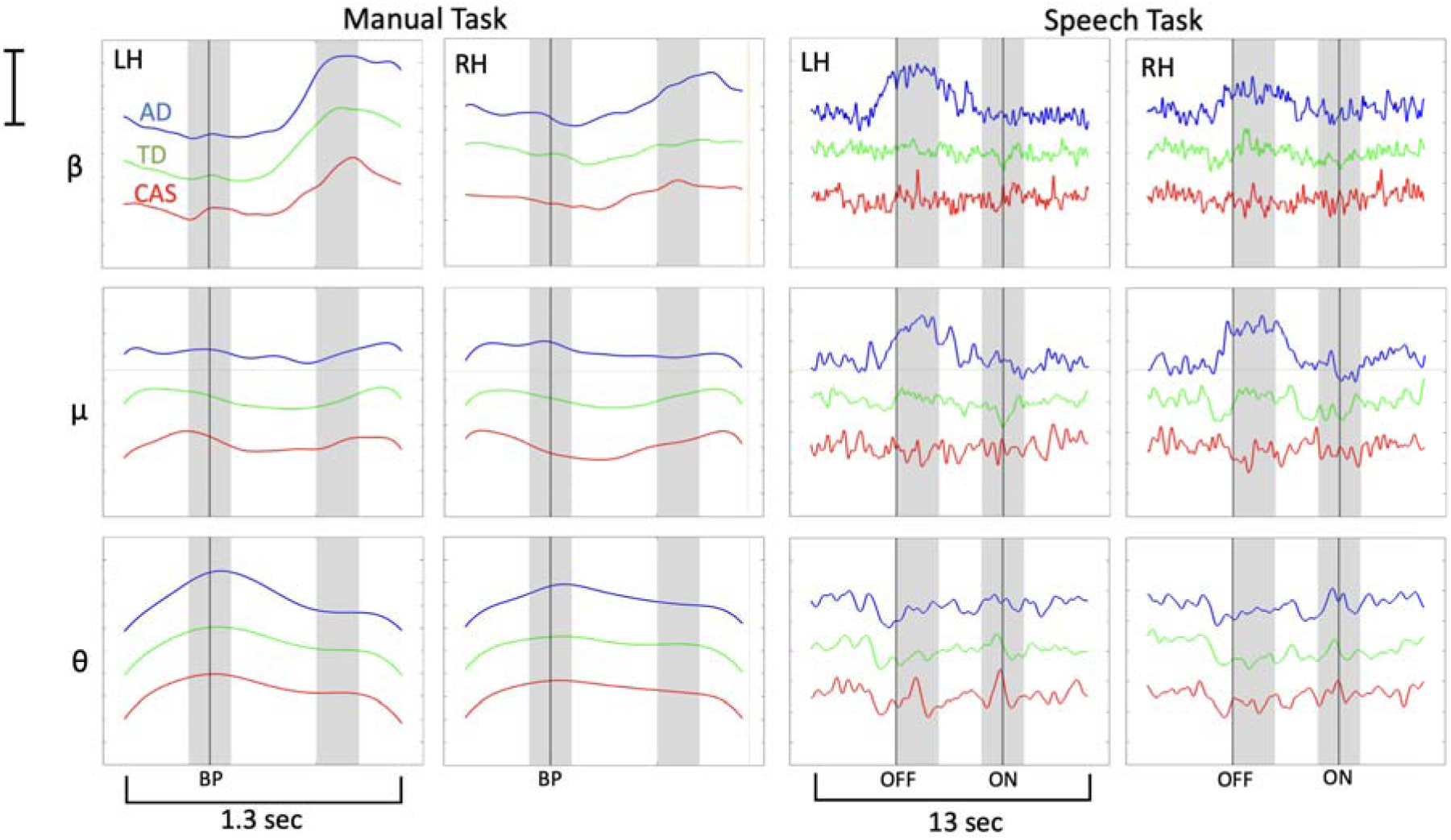
Group comparison of frequency-averaged time series showing beta-, mu- and theta-band responses elicited by manual and speech tasks in right and left hemispheres. Beginning and end of time series are truncated to remove edge artifacts from Fourier decomposition (see Figure 3). Shaded regions show ERS and ERD time-windows. LH = left hemisphere, RH = right hemisphere, BP = button press onset, OFF = speech trial set offset, ON = speech trial set onset. Scale bar indicates 25% change from baseline.

##### Response magnitude (within hemisphere)

Group comparisons of median speech response magnitudes are summarised in Table 4 and Figure 5. Nonparametric Kruskal-Wallis one-way ANOVAs indicated significant group differences in beta-band response magnitude in the left hemisphere (AD *Mdn* = 23.14, TD *Mdn* = 6.20, CAS *Mdn* = 0.00, χ^2^ (2,33) = 15, p < .001) but not for the right hemisphere (AD *Mdn* = 4.01, TD *Mdn* = 8.83, CAS *Mdn* = 4.27, χ^2^ (2,33) = 2.64, *p* = .267). Planned comparisons for the left hemisphere confirmed significantly greater response magnitude in AD than TD (p < .001) but no significant difference between CAS and TD (*p* = .298).

**Figure 5.**
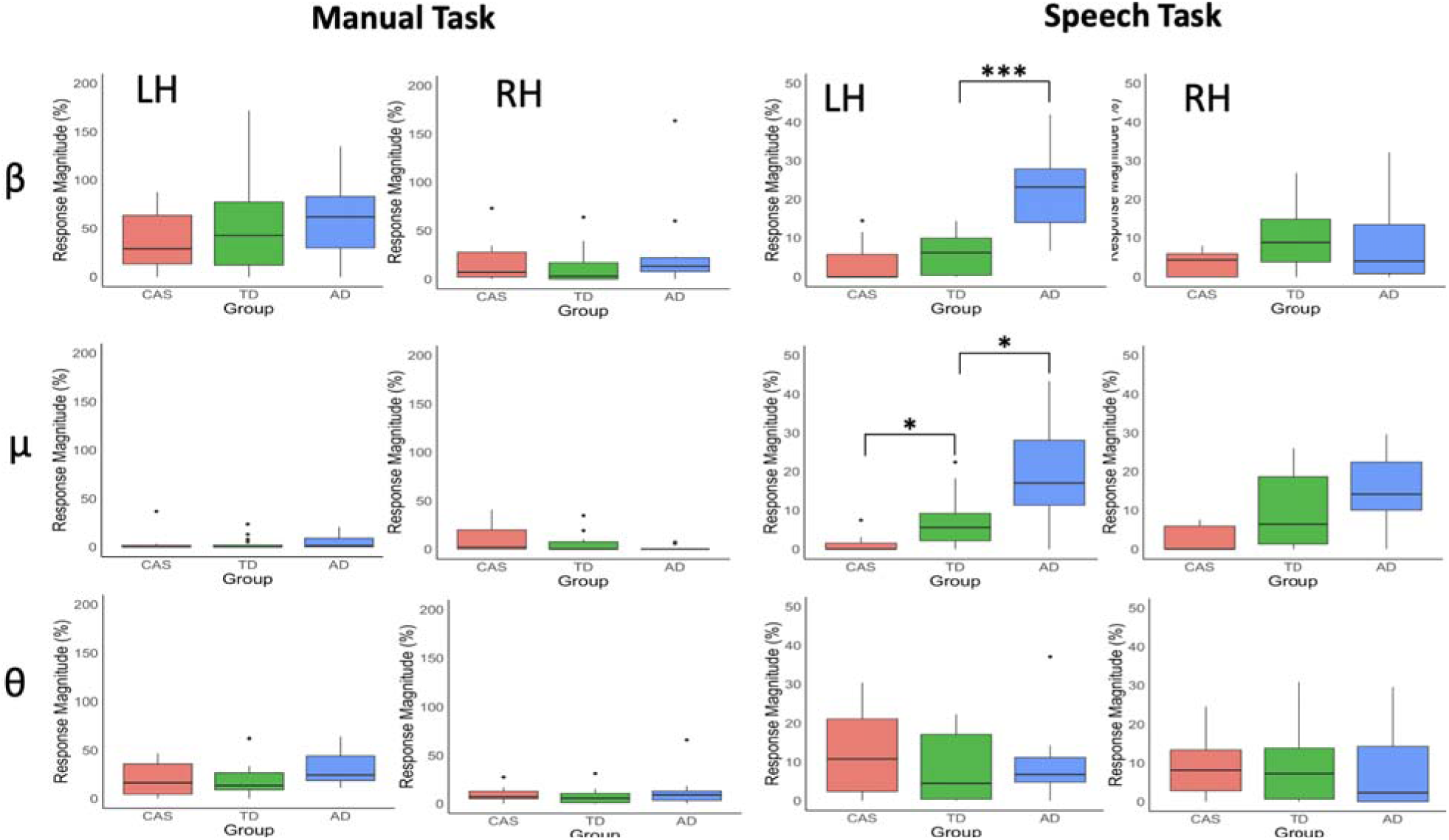
Group comparison of response magnitudes for beta, mu and theta bands elicited by manual and speech tasks in right and left hemispheres. Horizontal lines indicate median magnitude, boxes indicate interquartile range, whiskers indicate range. Note larger y-scale range for manual task than for speech task. *** = p <.001, * = p <.05.

**Table 4.**
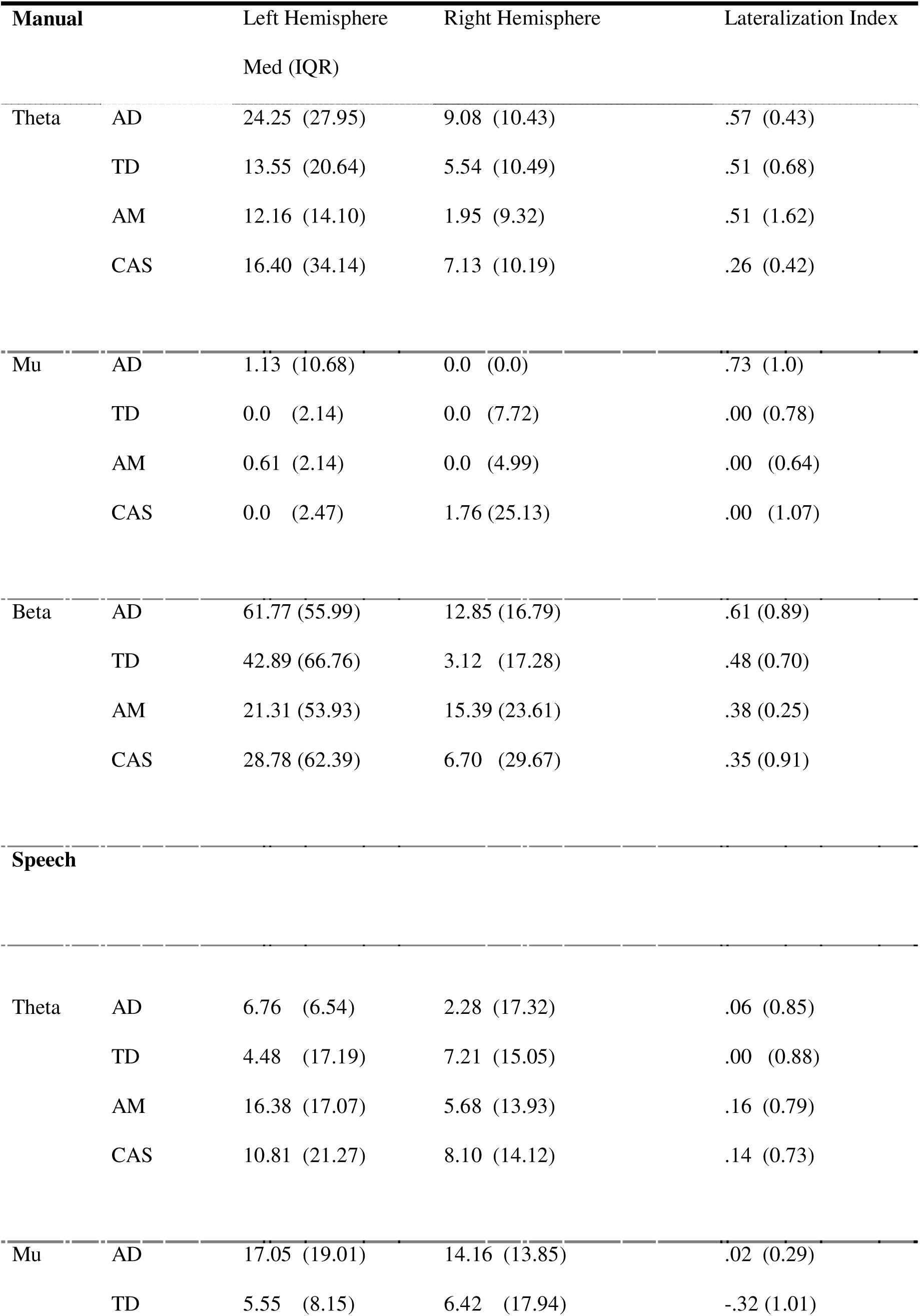
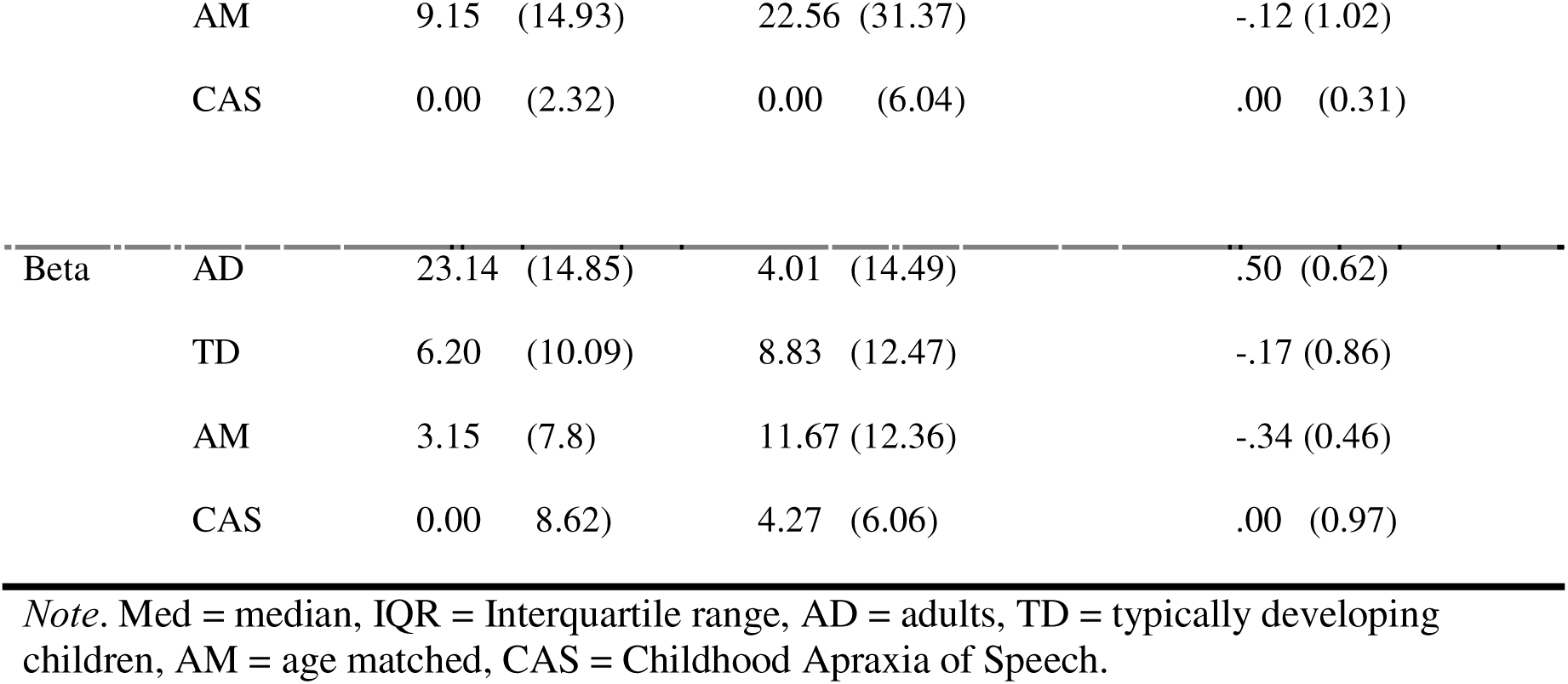
Response Magnitudes.

| **Manual** |  | Left Hemisphere | Right Hemisphere | Lateralization Index |
| --- | --- | --- | --- | --- |
|  |  | Med (IQR) |  |  |
| Theta | AD | 24.25 (27.95) | 9.08 (10.43) | .57 (0.43) |
|  | TD | 13.55 (20.64) | 5.54 (10.49) | .51 (0.68) |
|  | AM | 12.16 (14.10) | 1.95 (9.32) | .51 (1.62) |
|  | CAS | 16.40 (34.14) | 7.13 (10.19) | .26 (0.42) |
| Mu | AD | 1.13 (10.68) | 0.0 (0.0) | .73 (1.0) |
|  | TD | 0.0 (2.14) | 0.0 (7.72) | .00 (0.78) |
|  | AM | 0.61 (2.14) | 0.0 (4.99) | .00 (0.64) |
|  | CAS | 0.0 (2.47) | 1.76 (25.13) | .00 (1.07) |
| Beta | AD | 61.77 (55.99) | 12.85 (16.79) | .61 (0.89) |
|  | TD | 42.89 (66.76) | 3.12 (17.28) | .48 (0.70) |
|  | AM | 21.31 (53.93) | 15.39 (23.61) | .38 (0.25) |
|  | CAS | 28.78 (62.39) | 6.70 (29.67) | .35 (0.91) |
| **Speech** |  |  |  |  |
| Theta | AD | 6.76 (6.54) | 2.28 (17.32) | .06 (0.85) |
|  | TD | 4.48 (17.19) | 7.21 (15.05) | .00 (0.88) |
|  | AM | 16.38 (17.07) | 5.68 (13.93) | .16 (0.79) |
|  | CAS | 10.81 (21.27) | 8.10 (14.12) | .14 (0.73) |
| Mu | AD | 17.05 (19.01) | 14.16 (13.85) | .02 (0.29) |
|  | TD | 5.55 (8.15) | 6.42 (17.94) | -.32 (1.01) |
| Speech motor cortex function in CAS |  |  |  |  |
|  | AM | 9.15 (14.93) | 22.56 (31.37) | -.12 (1.02) |
|  | CAS | 0.00 (2.32) | 0.00 (6.04) | .00 (0.31) |
| Beta | AD | 23.14 (14.85) | 4.01 (14.49) | .50 (0.62) |
|  | TD | 6.20 (10.09) | 8.83 (12.47) | -.17 (0.86) |
|  | AM | 3.15 (7.8) | 11.67 (12.36) | -.34 (0.46) |
|  | CAS | 0.00 (8.62) | 4.27 (6.06) | .00 (0.97) |

*Note.* Med = median, IQR = Interquartile range, AD = adults, TD = typically developing children, AM = age matched, CAS = Childhood Apraxia of Speech.

Mu band responses in both hemispheres show an AD > TD > CAS group ordering of median magnitudes. Kruskal-Wallis one-way ANOVA confirmed a significant overall group difference in both the left (AD *Mdn* = 17.05, TD *Mdn* = 5.55, CAS *Mdn* = 0.00, χ^2^ (2,33) = 11.26, *p* = .004) and right hemispheres (AD *Mdn* = 14.16, TD *Mdn* = 6.42, CAS *Mdn* = 0.00). Within the left hemisphere, planned contrasts confirmed a significant group difference between AD and TD (*p* = .023) and between TD and CAS (*p* = .018). Further, the contrast between CAS and the AM subset of TD (*Mdn* = 22.56) was also significant (*p* = .017). In the right hemisphere, however, neither planned contrast achieved statistical significance (CAS vs TD, *p* = .078; TD vs AD, *p* = .280).

For the theta band, Kruskal-Wallis one-way ANOVAs showed no significant group differences in response magnitude in either hemisphere.

##### Hemispheric differences (within-group)

Wilcoxon signed rank tests showed significant hemispheric differences in beta-band responses for adults (LH *Mdn* = 23.14, RH *Mdn* = 4.01, *p* = .012), TD (LH *Mdn* = 6.20, RH *Mdn* = 8.83, *p* = .045), and the AM subset of TD (LH *Mdn* = 3.15, RH *Mdn* = 11.67, p = .031) but not for CAS (LH *Mdn* = 0.00, RH *Mdn* = 4.27, *p* = .875).

No significant hemisphere differences in mu or theta band response magnitudes were obtained for any of the three groups.

##### Lateralisation index (between group)

For assessment of between-group differences in hemispheric lateralization, LI (as described in Methods) was computed from response magnitudes in each hemisphere. Figure 6 (right panel) summarises LI for the speech task. Kruskal-Wallis one-way ANOVA indicated a significant group difference for beta lateralisation index (AD *Mdn* = .50, TD *Mdn* = −.17, CAS *Mdn* = .00). Wilcoxon ranksum planned contrasts confirmed AD group was significantly more left-lateralised than TD (*p* = .006), but there was no significant difference for TD and CAS groups (*p* = .652).

**Figure 6.**
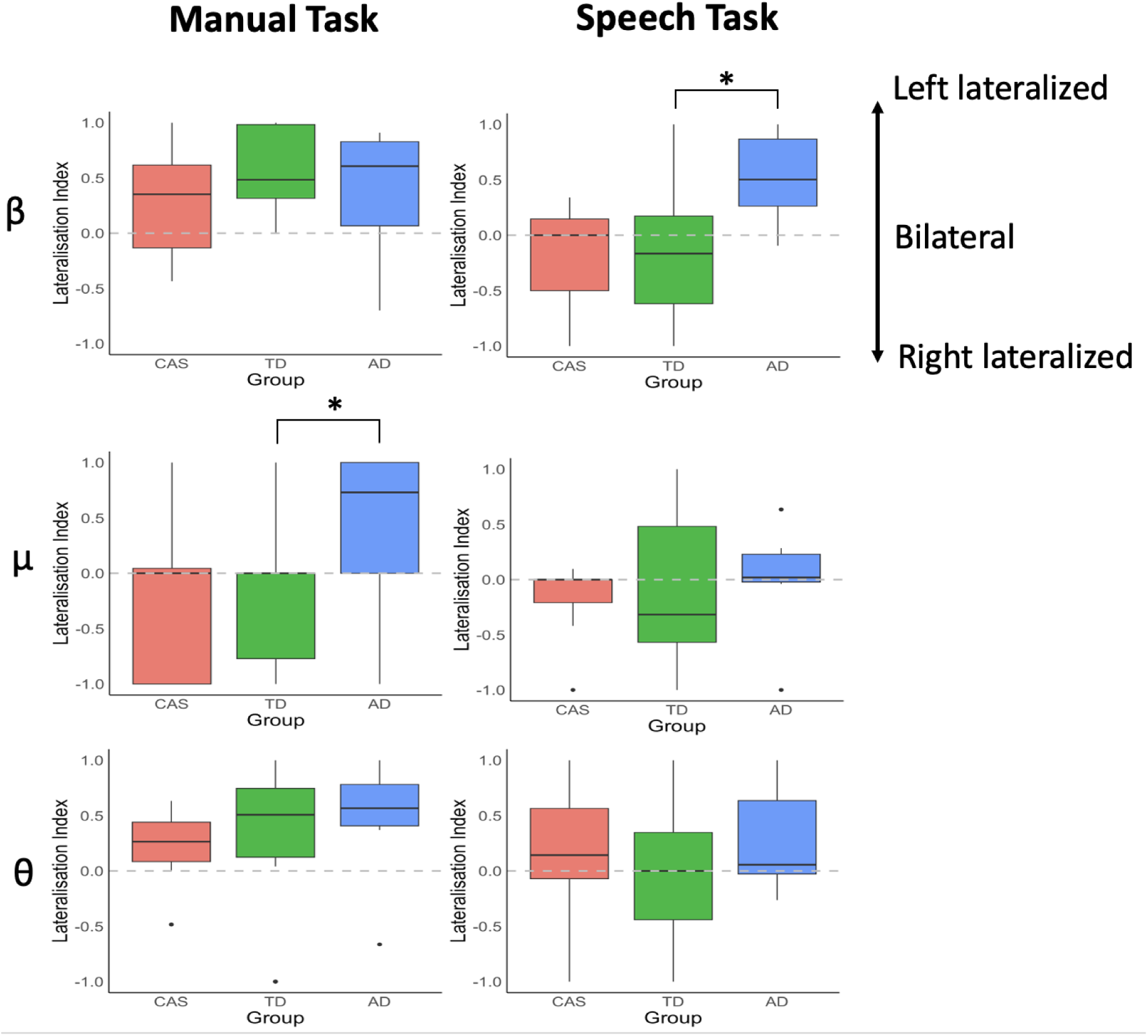
Group comparison of hemispheric lateralisation index for beta, mu and theta band responses to manual and speech task. * = p <.05.

ANOVAs showed no significant overall LI group differences for mu band (AD *Mdn* = .020, TD *Mdn* = −.32, CAS *Mdn* = .000, p = .40) or theta band (AD *Mdn* = .06, TD *Mdn* = .00, CAS *Mdn* = .14, *p* = .537).

##### Effects of age in TD children

Linear regression modelling indicated significant linear effects of age within the TD group only in the theta band response for the speech task (Supplementary Figure S1). For the speech task, theta response magnitude showed a significant linear increase with age in the right hemisphere (*F*(1,17) = .015, *R*^2^ = .300, p = .015). There was no significant effect of age in the left hemisphere (*p* = .375), and theta LI showed a significant decrease (i.e. increasing right lateralisation) with age (*F*(1,17) = 6.528, *R*^2^ = .277, *p* = .021). Overall, within the 7-16 yo range of the TD group, significant (albeit small) age effects were obtained only in theta band speech responses, with no indication of any (linear) maturational trajectory that could account for the prominent differences between the TD and adult mu/beta band speech responses. ***Summary of speech task results***. The three groups showed distinct patterns of brain activity for the speech task. While SAM beamformer clusters indicated activations in proximity to the hand task clusters near periRolandic sensorimotor cortex, adults were significantly more (left) lateralised than TD. We obtained no significant difference in overall hemispheric lateralisation between TD and CAS for any of the three frequency bands; however while TD showed smaller magnitudes of speech-related beta/mu responses than AD, these were virtually absent in the CAS children. A group ordering of response magnitudes was apparent for the left hemispheric mu-band response, which was significantly lower in magnitude relative to both the overall TD group and the subset of age-matched TD.

Relative to the adults, the child groups showed speech-elicited clusters that were deeper and more ventral. Group discrepancies in the precise locations of clusters are difficult to interpret from the present data. On the one hand, good between-group agreement of source locations for the manual task supports adequacy of source modelling for all groups: in this case all groups showed good agreement for *x* and *y* coordinates, while the *z* discrepancy is explainable from the more extensive cluster configuration in the adults. On the other hand, group discrepancies may be expected to arise from three sources in the present experiment: first, adult source modelling was based on individual MRI scans, while the child models were derived from template brains; second, the cluster-based analysis indicates both more extensive and stronger (larger magnitude) sources for the adults than for the child groups, a difference in signal-to-noise ratios that may affect model comparisons; and third, it is known that adaptive beamformers perform suboptimally in the case of correlated bilateral sources (as in the TD group), since linear dependencies between the neuronal source timeseries are utilised by these algorithms to minimize the output power (e.g., (Kuznetsova et al., 2021)).

Overall, these considerations preclude any clear inferences concerning adult and child group differences in the precise locations of speech-related sources. Within the stated limitations, the results of the statistical cluster-based analyses suggest the following: for adults, a differential SAM beamformer effect (p < .05) corresponds to a single cluster in the observed data at a spatial location in the precentral gyrus of the left hemisphere immediately inferior to the known anatomic location of the hand region of sensorimotor cortices, and co-extensive with the region of the medial frontal gyrus that is immediately anterior to this middle region of the pre-central gyrus; in contrast, for the TD group, the differential SAM beamformer effects (p < .05) correspond to bilateral clusters in both cerebral hemispheres, at relatively lateralised locations on a mid-line (in the sagittal plane) roughly corresponding to the location of the central sulcus/ pre- and post-central gyri in both hemispheres; while for the CAS group the significant beamformer effects correspond to a single cluster in the right hemisphere corresponding roughly to the location of the right hemisphere cluster of the TD group.

#### b. Manual task

##### Source reconstruction

Group mean and corresponding statistical maps of SAM-beamformer images for the manual condition are shown in Figure 2 (left panels). Consistent with the previous literature, all groups showed maximal responses in the regions of the left precentral gyrus (contralateral to the right-handed button press) with centre coordinates (Table 3) located at or near the hand region of the hand knob of the precentral gyrus. Also consistent with previous work, all groups also showed mirrored right hemisphere activations in homologous regions of the right hemisphere motor cortex (not shown), although these clusters were smaller in magnitude and did not reach statistical significance for any of the groups.

##### Virtual sensors

Group mean virtual sensor time-frequency plots were generated from the button press coordinates listed in Table 3 and are plotted in the left panels of Figure 3. The following features can be observed in the plots for all three groups: (1) Beta-band (circa 13-30 Hz) desynchronisation beginning about 300 ms before the button press and persisting for about 300-400 ms after the button press; (2) Beta-band synchronisation (beta “rebound”) beginning about 500-600 ms after the button press and persisting for 300-400 ms; theta-band (circa 3-7 Hz) synchronisation beginning about 200-300 ms prior to the button press and persisting until 500-600 ms post-button press. Comparable but lower magnitude spectral perturbations are also evident in the right hemisphere spectrograms. All of these temporal-spectral features are known characteristics of neuromagnetic brain responses in a self-paced button press task for both adults and children and have been described in previous publications (e.g. Cheyne et al., 2014; Johnson et al., 2020; Fung et al., 2022). Group comparisons of frequency-band averaged time series (Figure 4) further confirm that the three groups exhibit entirely comparable brain activity during manual movements.

##### Response magnitude (within hemisphere)

Further statistical analyses of event-related spectral perturbations were carried out using the response magnitude metric (see Methods). Group comparisons of median response magnitudes are summarised in Table 4 and Figure 5. Nonparametric Kruskal-Wallis one-way ANOVAs showed no significant group differences in response magnitude within either hemisphere, for any of the three frequency bands.

##### Hemispheric differences (within-group)

Hemispheric differences were assessed within each group using nonparametric Wilcoxon signed rank tests (2-sided). Results showed that beta-band response magnitudes were significantly higher in the left hemisphere for AD (*N* = 11, L *Mdn* = 61.77, R *Mdn* = 12.85, *W* = 34, *p* = .023), and TD (*N*= 19, L *Mdn* = 42.49, R *Mdn* = 3.12, *W* = 153, *p* < 0.001). While median left hemisphere response magnitudes were also greater in the CAS group, the hemisphere difference did not reach statistical significance (N = 7, L *Mdn* = 28.78, R *Mdn* = 6.70, *W* = 15, *p* = .438), but did for the AM subset of TD (*N* = 7, L *Mdn* = 21.31, R *Mdn* = 15.39, *W* = 28, *p* = .016).

In the mu band no significant hemispheric differences in response magnitude were obtained for any group, while in the theta band a significant hemispheric difference in response magnitude were obtained only for the TD group (*N* = 19, L *Mdn* = 13.55, R *Mdn* = 5.54, *z* = 2.656, *W* = 161, *p* = .008).

##### Lateralisation index (between group)

Nonparametric Kruskal-Wallis one-way ANOVA showed no significant group differences for beta-band LI, with all groups showing a left lateralised profile (AD *Mdn* = .61, TD *Mdn* = .48, CAS *Mdn* = .35, χ^2^(2,33) = 2.67, *p* = .263). Kruskal-Wallis one-way ANOVA indicated a significant group difference for mu LI, with only adults showing a left-lateralised response (AD *Mdn* = .73, TD *Mdn* = .00, CAS *Mdn* = .01, χ^2^ (2,33) = 6.44, *p* = .040). Wilcoxon ranksum planned comparisons confirmed significantly greater left lateralisation for AD compared to TD groups (*p* = .013), but there was no significant difference between TD and CAS groups (*p* = .952). No significant group differences were obtained for theta band LI and all groups showed a left lateralised profile (AD *Mdn* = .57, TD *Mdn* = .51, CAS *Mdn* = .26, χ^2^ (2,33) = 2.94, *p* = .230).

##### Summary of manual task results

In contrast to the results of the speech task, all three groups showed entirely comparable patterns of brain activity during the manual movement task, with SAM beamformer clusters indicating activation at or near the contralateral hand region of sensorimotor cortex, and with smaller (and statistically nonsignificant) activations in homologous regions of the right hemisphere. Spectro-temporal patterns computed from these regions were prominently characterised by a robust pattern of event-related beta-band desynchronisation prior to and following the button press movement; and followed after about 500-600 ms by a “rebound” beta-band synchronisation. Within the theta band the spectral response followed a comparable time-course with opposite polarity (i.e. theta ERS during mu/beta ERD, and theta ERD during mu/beta ERS). As the only significant group difference, adults showed greater left hemisphere lateralisation of mu-band activity than TD, although mu-band activity was notably small in magnitude in all groups and both hemispheres. Overall, we conclude that the cluster-based permutation analyses show a significant differential SAM beamformer effect for all three groups, with clusters in the observed data at a spatial location corresponding to the known anatomic location of the hand region of sensorimotor cortex in the left hemisphere, contralateral to the right index finger used for the button press task. Further, virtual sensor time-spectrograms generated from the cluster centre locations for each group are entirely representative of and in accord with known and well-characterised neuromagnetic brain responses in a self-paced button press task (e.g. Cheyne et al., 2014; Fung et al., 2022; Johnson et al., 2020).

## Discussion

The present results provide the first evidence for dysfunction in the neurophysiological generators of the mu-band speech motor rhythm in a sample of children with clinically diagnosed CAS. Our analyses specify a brain origin of this speech rhythm in the left cerebral hemisphere, within or near pre-Rolandic motor areas crucial for the planning and control of speech and oromotor movements. We further refine this anatomical specification by demonstrating the proximity of speech-elicited beamformer maps to clusters associated with manual button-press responses known to originate in the hand area of the peri-Rolandic sensorimotor cortex. Taken together, these results provide clear functional-anatomical delineation of neural activity as a candidate biomarker of childhood speech apraxia.

### Apraxia of speech

CAS is a clinical subtype of AOS and is differentiated from acquired forms of AOS by a developmental aetiology and an absence of frank lesions from acquired brain injuries. Despite differences in clinical presentation due to the aetiological dissimilarities, the two subtypes share many apraxic features likely to be attributable to commonalities in their causal neuropathologies (Miller & Guenther, 2021). While differential diagnosis of CAS is an area of active and ongoing research (Murray et al., 2015), features identical to those used for acquired AOS – including problems with syllable segregation, syllable stress, and speech sound distortions – have been proposed and are increasingly used for diagnosis of CAS (Miller & Guenther, 2021; Murray et al., 2015; Shriberg et al., 2009, 2017; Yoss & Darley, 1974). Psycholinguistic and motor control models have long attributed these apraxic features to disruptions of motor programming due to dysfunction at the interface between otherwise intact phonological and motor systems, an interface wherein abstract linguistic codes are transformed into movement commands for interpretation by the motor system (ASHA, 2007).

The current results dovetail neatly with evidence implicating the left precentral gyrus in acquired AOS. In particular, cases of “pure” acquired AOS (i.e. AOS features without other speech-language deficit) have consistently been associated with lesions of the left precentral gyrus (Basilakos et al., 2015; Graff-Radford et al., 2014; Itabashi et al., 2016; Moser et al., 2016; Robin et al., 2008; Takakura et al., 2019). Further evidence for the role of the left precentral gyrus is provided by reports that neurostimulation of this region results in transient disruptions of speech with the main features of AOS (Duffau et al., 2003; Tandon et al., 2002). Finally, neurocomputational lesion modelling studies (Guenther, 2016; Miller & Guenther, 2021) have reported that AOS speech symptomology can only be produced in these models through damage to a putative speech sound map located in the left ventral precentral gyrus and surrounding portions of posterior inferior frontal gyrus and anterior insula. In contrast, lesions to most other components of the model resulted in dysarthria rather than AOS. According to this computational model, AOS symptomology is dependent specifically on damage to the speech sound map in the *left* ventral precentral gyrus which is tasked with feedforward control of speech motor programs (while its right hemisphere homologue is proposed to be specialised for *feedback* motor control of corrective motor commands). (Guenther, 2016).

### Speech-elicited brain activities

Relative to the extended and distributed patterns of activations that are typically obtained with lexical speech production tasks (Agarwal et al., 2019; Munding et al., 2016), our results show spatially-restricted clusters associated with peri-Rolandic regions of sensorimotor cortex. These loci and their relative focality concur with the fMRI results using a comparable reiterated speech task reported by Riecker et al. (2000). These authors suggested that overlearned, automatic, and repetitive utterances can be effectively “chunked” at a planning level in a manner that places far fewer demands on neural resources than are required for single “individualised” speech gestures. In addition, the lack of lexical content in the nonword utterances removes the need for any access to higher language centres for syntactic processing or lexical retrieval.

Our results and those of Riecker et al. (2000) are entirely supportive of an emerging neuroscientific consensus which assigns a central role in speech motor planning and coordination to speech motor regions of the precentral gyrus and immediately adjacent regions of the prefrontal cortex (Glasser et al., 2016; Gordon et al., 2023; Jensen et al., 2023; Silva et al., 2022; Willett et al., 2023). This new conceptualization is exemplified in the recent “dual motor system” model of expressive speech processing (Hickok et al., 2023), which posits a hierarchical control system consisting of dorsal and ventral portions of the precentral gyrus and the posterior portion of the middle frontal gyrus.

### Limitations and future directions

The major limitation of our CAS data is a small group size which is common for published studies of this rare disorder (Table 1). Offsetting this are a number of important methodological and analytical strengths: The CAS phenotype was rigorously characterised (Chenausky & Tager-Flusberg, 2022) with a comprehensive battery of clinical tests and diagnosed independently on the basis of overt speech behaviours by two experienced speech-language pathologists; the locus of the speech cluster in the CAS group is in accord with that of the right hemisphere cluster of TD children; and the neurophysiological responses of all seven individuals show absence or very low magnitudes of left hemisphere speech-related mu-band activity that is a prominent feature of both the TD and AD groups (Supplementary Figure S1). Therefore, despite the small sample size, group mu-band magnitude differences were statistically significant for both the CAS-TD group comparison and the CAS-AM comparison. These considerations are indicative of a robust and sensitive neurophysiological marker which has substantial potential for application in studies of typical and apraxic speech development.

Our MEG results for the manual task indicate developmentally typical hand motor cortex function in the CAS group, at least for the simple movements required in this task. This provides an intriguing contrast to the neurophysiological anomalies associated with the speech task, particularly in light of our behavioural data indicating significant deficits for non-speech motor skills. Fine and gross non-speech motor deficits suggestive of developmental coordination disorder (DCD) are estimated to occur in some 50%-80% of children with CAS, (Duchow et al., 2019; Iuzzini-Seigel, 2021; Iuzzini-Seigel et al., 2022;), a co-occurrence that has invited considerable speculation about shared mechanisms within the central motor system (And & Dodd, 1996; Duchow et al., 2019; Iuzzini-Seigel, 2021; Iuzzini-Seigel et al., 2022; Knežević, 2019; McCabe et al., 1998). While the present data from a simple button press task are not readily comparable to to the sequential and coordinated articulatory movements required in the speech task, the basic experimental framework can be readily adapted for future studies wherein the relative functioning of speech and hand motor cortices can be probed using tasks where movement complexity is better equated (see for e.g. (De Nil et al., 2021).

## Acknowledgements

This work was supported by a Waterloo Foundation Child Development Fund Research Grant (Ref. No. 2532-4758) and a Discovery Project Grant (DP170102407) from the Australian Research Council.

Ioanna Anastasopoulou’s current affiliation is Hospital for Sick Children Research Institute, Toronto, Ontario M5G 0A4, Canada.

## Supplementary Materials

**Supplementary Table S1.** Individual test results for child groups.

| Appendix 1 Table S1: Individual test results for child groups |  |  |  |  |  |  |  |  |  |  |  |  |
| --- | --- | --- | --- | --- | --- | --- | --- | --- | --- | --- | --- | --- |
|  | TD |  |  |  |  |  |  |  |  |  |  |  |
| Age Match | Participant number | Age | Sex | Expressive and Receptive Language (CELF-5) | Articulation in Words (GFTA-3) | Articulation in Sentences (GFTA-3) | VMPAC GMC % | VMPAC FOMC % | VMPAC SEQN % | VMPAC CSP % | VMPAC SPC % | MACB-2 SS Total |
| 1 | 1158 | 8.72 | 1 | 9 | 100 | 113 | 100 | 99.25 | 100 | 100 | 100 | 8 |
| 2 | 1160 | 8.53 | 1 | 11 | 110 | 114 | 100 | 99.63 | 100 | 100 | 100 | 14 |
| 3 | 1145 | 9.55 | 0 | 17 | 106 | 109 | 100 | 98.88 | 100 | 100 | 100 | 12 |
| 4 | 1146 | 8.35 | 0 | 13 | 71 | 89 | 100 | 99.25 | 100 | 100 | 100 | 12 |
| 5 | 1171 | 7.47 | 0 | 16 | 103 | 121 | 100 | 100 | 100 | 100 | 100 | 9 |
| 6 | 1138 | 8.15 | 1 | 7 | 111 | 115 | 100 | 95.9 | 93.48 | 100 | 100 | 9 |
| 7 | 1156 | 12.01 | 1 | 25 | 105 | 109 | 100 | 100 | 100 | 100 | 100 | 10 |
| 8 | 1140 | 12.89 | 1 | 25 | 105 | 109 | 100 | 98.51 | 100 | 100 | 100 | 14 |
| 9 | 1141 | 16.73 | 1 | 26 | 102 | 107 | 100 | 100 | 100 | 100 | 100 | 14 |
| 10 | 1148 | 13.61 | 0 | 25 | 103 | 107 | 100 | 100 | 100 | 100 | 100 | 10 |
| 11 | 1149 | 11.55 | 0 | 29 | 104 | 107 | 100 | 100 | 100 | 100 | 100 | 9 |
| 12 | 1151 | 11.97 | 1 | 28 | 105 | 109 | 100 | 99.25 | 100 | 100 | 100 | 12 |
| 13 | 1166 | 12.86 | 0 | 27 | 104 | 107 | 100 | 100 | 100 | 100 | 100 | 15 |
| 14 | 1169 | 14.64 | 1 | 25 | 103 | 108 | 100 | 100 | 100 | 100 | 100 | 8 |
| 15 | 1172 | 12.43 | 1 | 23 | 105 | 109 | 100 | 98.13 | 100 | 100 | 100 | 13 |
| 16 | 1178 | 11.06 | 1 | 15 | 89 | 80 | 100 | 100 | 100 | 100 | 100 | 11 |
| 17 | 1147 | 9.37 | 0 | 27 | 106 | 109 | 100 | 98.88 | 100 | 100 | 100 | 8 |
| 18 | 1163 | 9.8 | 0 | 19 | 108 | 111 | 100 | 100 | 100 | 100 | 100 | 10 |
| 19 | 1165 | 9.02 | 1 | 23 | 111 | 108 | 100 | 99.63 | 100 | 100 | 100 | 9 |
|  | CAS |  |  |  |  |  |  |  |  |  |  |  |
| 1 | 1154 | 7.98 | 1 | 12 | 59 | 56 | 85 | 75.75 | 91.3 | 86.67 | 57.14 | 6 |
| 2 | 1155 | 7.86 | 1 | 8 | 64 | 69 | 65 | 70.52 | 95.65 | 86.67 | 57.14 | 4 |
| 3 | 1164 | 9.67 | 1 | 11 | 40 | 42 | 70 | 66.69 | 76.09 | 82.22 | 42.86 | 7 |
| 4 | 1168 | 7.52 | 1 | 4 | 40 | 41 | 85 | 64.55 | 63.04 | 66.67 | 28.57 | 9 |
| 5 | 1170 | 6.79 | 1 | 5 | 66 | 85 | 100 | 84.33 | 89.13 | 75.56 | 71.43 | 11 |
| 6 | 1174 | 7.22 | 1 | 7 | 68 | 73 | 90 | 82.09 | 78.26 | 77.78 | 57.14 | 8 |
| 7 | 1181 | 12.8 | 1 | 26 | 79 | 75 | 95 | 86.57 | 97.83 | 84.44 | 85.71 | 7 |

**Supplementary figure S1.**
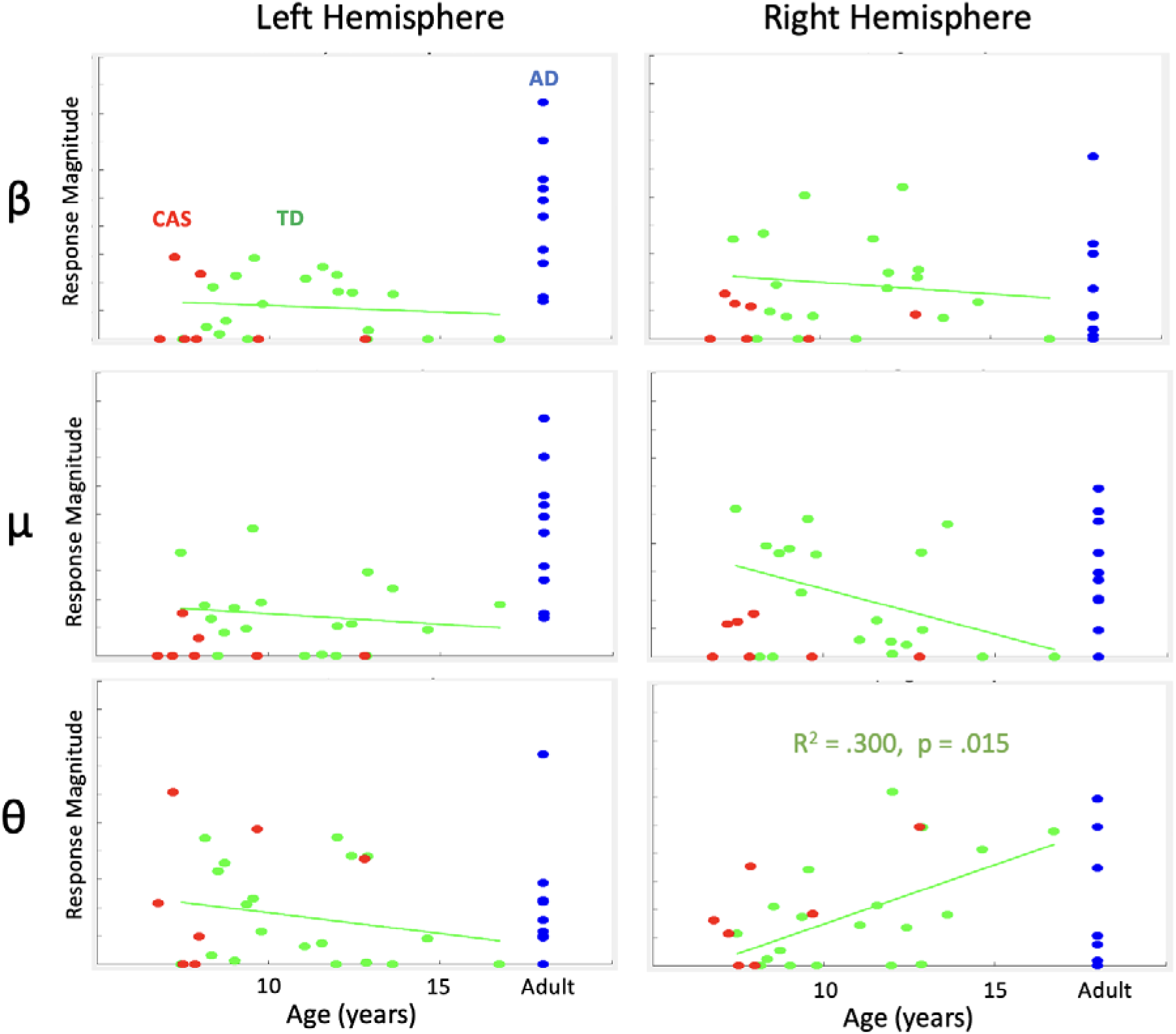
Individual response magnitudes for speech task for each group and hemisphere. TD and CAS group data points only are plotted against age. Linear regression line is plotted for TD group only.

